# A methylome-derived m^6^-dAMP trigger assembles a PUA-Cal-HAD immune filament that depletes dNTPs to abort phage infection

**DOI:** 10.64898/2026.01.15.699771

**Authors:** Zhiying Zhang, Yi Wu, Yan-Jiun Lee, Gianlucca G. Nicastro, Arpita Chakravarti, Hui Fang, Daniela Taverner, Peter Weigele, L. Aravind, Dinshaw J. Patel, Franklin L. Nobrega

**Author notes:** Shared first authors (Zhiying Zhang, Yi Wu).

## Abstract

Bacteria must distinguish phage attack from normal homeostatic processes, yet the danger signals that trigger many defence systems remain unknown. Here, we show that a PUA-Calcineurin-CE–HAD module from *Escherichia coli* ECOR28 confers broad anti-phage protection by binding Dam-methylated deoxyadenosine monophosphate (m^6^-dAMP) generated during phage-induced chromosome degradation. Ligand binding converts a preassembled PUA-Calcineurin-CE hexamer loaded with six HAD phosphatases into a polymerising filament. The filament acts as a high-flux dNTP sink through a two-enzyme cascade: HAD first dephosphorylates dATP to dADP, and Calcineurin-CE then converts dADP to dAMP. dNTP collapse halts phage replication and enforces abortive infection. Multiple mobile-element DNA mimic proteins block filament assembly, revealing a direct phage counter-defence. More broadly, our findings extend a conserved, cross-kingdom paradigm of immune filament assembly to nucleotide-depletion antiviral defence and suggest modified-nucleotide sensing by related PUA-Calcineurin-CE modules as a widespread, underappreciated bacterial strategy.

## Introduction

Bacteria are locked in an arms race with bacteriophages and deploy a battery of defence systems to survive viral infection. Recent computational and experimental surveys have revealed an unexpectedly large repertoire of such systems^1,2^. Many newly discovered loci confer anti-phage protection, yet their molecular mechanisms remain only partially understood^1–7^. Others are currently recognised as candidate defence systems based on sequence and genomic context^8–10^. One such candidate is a two-gene module encoding a PseudoUridine synthase and Archaeosine transglycosylase (PUA)-like sensor fused to a Calcineurin-conflict effector (CE) phosphoesterase (Cal), together with a partner haloacid dehalogenase (HAD) phosphatase (hereinafter PUA-Cal-HAD) ^8^.

Calcineurin-CE is a recently identified family within the calcineurin-like superfamily of metallophosphoesterases, predicted to function as an effector in diverse prokaryotic anti-phage immune systems^8^. Contextual analysis suggested that Calcineurin-CE domains act by targeting cellular nucleotide pools, thereby restricting phage replication through depletion of essential metabolites. In the two-gene module considered here, Calcineurin-CE is fused to an N-terminal PUA-like domain, raising the possibility that nucleotide depletion is coupled to a ligand-sensing input.

PUA-like domains were first described as RNA-binding modules in RNA-modifying enzymes, where they contribute to the recognition of structured and chemically modified RNA^11–13^. Subsequent structural and biochemical work has identified PUA-like folds across a wide range of DNA- and RNA-associated proteins^14–16^. The fold typically contains a compact binding pocket in which nucleobases are cradled by aromatic and charged residues, enabling binding to both canonical and chemically modified nucleotides^15,16^. Consistent with this biochemistry, PUA-like domains are expanded in conflict-associated systems that are predicted to sense epigenetic marks in the form of (hyper)-modified nucleobases, as cues for self-nonself discrimination or infection-associated states^8,17,18^.

The second gene in this module encodes a HAD superfamily phosphatase. HAD enzymes are Mg^2+^-dependent phosphatases that typically hydrolyse phosphomonoesters, including metabolites derived from sugars, nucleotides and cofactors^19,20^. On the basis of domain composition and genomic coupling, it was proposed that the antiviral mechanism of these loci involves concerted phosphohydrolase activities of Calcineurin-CE and HAD acting on nucleotide pools, with the PUA-like domain enabling regulated deployment of the effector response^8^. However, both the antiviral activity and its molecular mechanism have remained untested.

Here we characterised a representative of PUA-Cal-HAD from *Escherichia coli* ECOR28 and defined its defence activity and mode of action. We show that PUA-Cal-HAD confers broad anti-phage protection and requires the PUA-like sensor, the Calcineurin-CE and HAD effectors, together with their ligand-binding and catalytic motifs. Structural analyses reveal that the PUA-Cal fusion forms a cation-stabilised hexamer that recruits six HAD molecules at Calcineurin-CE interfaces, preassembling a large effector complex. We further show that phage-induced degradation of Dam-methylated host DNA generates methylated adenine nucleotides that bind the PUA-like domain and trigger polymerisation into filaments. Upon activation, HAD first dephosphorylates dNTPs to dNDPs, and Calcineurin-CE then converts dNDPs into dNMPs, collapsing cellular dNTP pools. This coordinated two-enzyme effector mechanism arrests phage replication and host growth, imposing abortive infection and revealing a methylome-derived danger signal that activates filament-driven nucleotide-depletion immunity. Finally, we show that multiple DNA mimic proteins can antagonise this defence by binding to the PUA-like domain and preventing higher-order assembly.

## Results

### Computational analysis and discovery of novel systems utilising PUA-like, Calcineurin-CE and HAD domains

The initial analysis of the Calcineurin-CE domains^8^ has led to the identification of the PUA-Cal-HAD module. To better constrain our functional predictions regarding its individual components, we utilised the principle that biological conflict systems have evolved combinatorial diversity through the tendency to mix-and-match effector and regulatory domains^21^. Thus, by studying the various contexts in which these domains occur, it is possible to computationally construct domain-architectural and gene-neighbourhood networks and algorithmically detect communities in them to predict the functions of poorly understood domains. Accordingly, we initiated sequence-profile searches with PUA-like, Calcineurin-CE and HAD domains to delineate the families of these domains found in these immune systems within their respective superfamilies and extracted their domain architectural and conserved-gene neighbourhood contexts.

Consistent with the prior study on Calcineurin-CE domains^8^, we invariably found them to be located in the effector context of diverse predicted anti-selfish element systems, suggesting that they primarily function as an effector in these systems (**Figure 1, Figure S1A,** genomic coordinates in **Table S1**). In addition to being fused to the Calcineurin-CE domain, we found the PUA-like domain to be fused to several other effector domains with unrelated catalytic activities (**Figure 1, Figure S1A,B**), such as the HEPN (endoRNase), DrHyd, TIR (both likely targeting NAD^+^), the HNH restriction endoDNase and the McrB AAA+ NTPase domain from McrBC-like restriction systems (the last two target DNA). These associations indicated that the PUA-like domain is likely to recruit a diverse array of effectors that either act by targeting invader nucleic acids or cellular biomolecules to respectively restrict infection or induce abortive infections through dormancy or cell death. Moreover, exemplars of this family of PUA-like domains were also found embedded in Restriction-Modification (R-M) operons with N^6^-adenine methylases (MTases). Further, this family of PUA-like domains associated with Calcineurin or DrHyd effectors is also linked to N^6^-adenine methylases independently of restriction enzymes, and operons for synthesis of modified bases (**Figure 1, Figure S1**). One of the latter codes for the enzyme adenylosuccinate synthetase, involved in the synthesis of N^6^-succinylated AMP. These observations, taken together with the known role of other PUA-like domains in binding modified nucleic acids and conservation of key residues in the nucleobase binding cleft (**Figure S1**), indicate that the PUA-like domain in these systems is likely to bind a modified nucleobase, which might include N^6^-methylated or hypermodified adenines.

**Figure 1.**
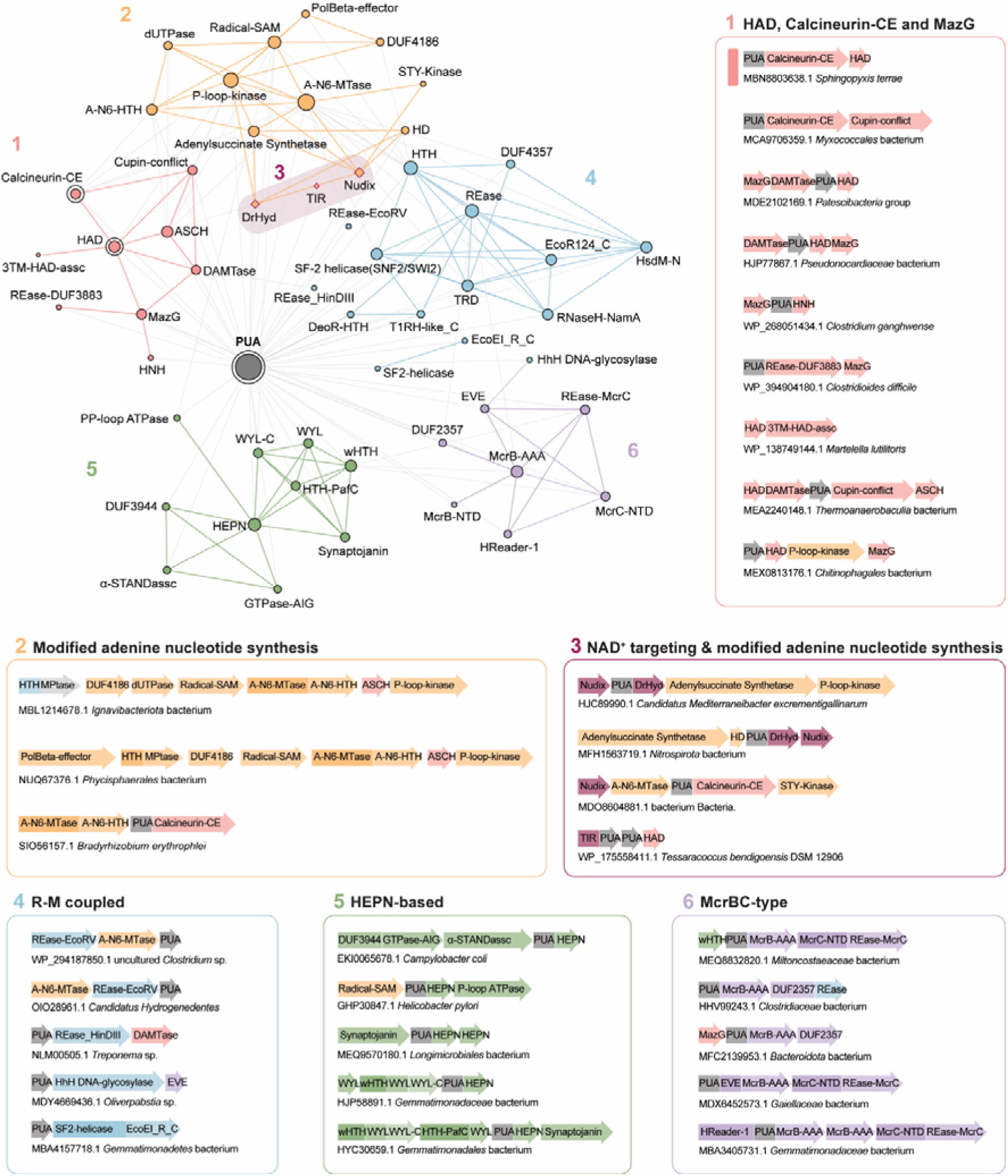
Contextual network of predicted anti-phage systems featuring a PUA-like domain. The key nodes interrogated in this study – PUA-like, HAD, and Calcineurin-CE protein domains – are shown with double circles. Nodes represent protein domains, and edges represent their co-occurrence in the form of domain fusions within the same protein or adjacency within the same gene operon. Nodes and edges are coloured according to inferred functional categories and numbered per group. Edges connecting nodes within the same dense subgraph (community) are coloured according to the corresponding theme, whereas edges linking nodes from different communities are shown in grey. Node size is proportional to degree, reflecting the number of unique genomic architectures. The purple box highlights a subgroup of NAD^+^-targeting systems within the larger group involving modified adenine nucleotide synthesis domains. Selected conserved gene neighbourhoods are shown on the right side and below the network. A pink band highlights the minimal tripartite architecture composed of PUA-like, HAD, and Calcineurin-CE domains. A detailed set of gene neighbourhoods used to construct the network are provided in **Figure S1** and the genomic coordinates are provided in **Table S1**. **(A)** Representative genomic contexts for the PUA-like domain family under consideration, grouped by shared functional themes. Genes are depicted as box arrows, with arrowheads indicating the 3′ ends of genes. Genes encoding multi-domain proteins are segmented into labelled regions corresponding to individual domains. Domains associated with modified nucleotide synthesis are highlighted in orange, whereas domains associated with NAD⁺ targeting are highlighted in magenta. **(B)** Multiple alignment of the PUA-like domain family considered in this study. The asterisks above the residues mark the positions predicted to be involved in recognition of modified nucleobase. The alignment is coloured according to the 90% consensus.

The HAD phosphatase domain found in these systems shows a strong association with Calcineurin-CE as the predicted effector. Notably, it is mostly absent when any of the other above-noted catalytically diverse effector domains are coupled to the PUA-like domain in lieu of the Calcineurin-CE domain (**Figure 1, Figure S1**). However, remarkably, there is a relatively rare set of systems wherein a MazG-like phosphoesterase replaces the Calcineurin-CE domain; nevertheless, these retain both the PUA and HAD components. Taken together with the previously reported activity of MazG-like proteins^22^, these observations suggested that both MazG and Calcineurin-CE potentially evince their effector action by targeting nucleotide triphosphates, with the HAD phosphate activity being required to augment their effector action.

Based on these predictions, we inferred that the protein with the PUA-like and Calcineurin-CE and that with the HAD domain form a complex of unknown stoichiometry, which remains negatively regulated under default conditions. However, upon the sensing of a modified nucleobase by the PUA-like domain (a likely infection signal), it undergoes a conformational change unmasking its effector activities.

### PUA-Cal-HAD is a tripartite anti-phage system that aborts infection

To test our hypothesis, we first assessed whether a PUA-Cal-HAD variant from *E. coli* ECOR28 (82599-85081 bp, Genbank accession no. NZ_LYAI01000252, Figure 2A) provides anti-phage protection by expressing it under its native promoters in *E. coli* BL21-AI. We challenged these cells with a panel of phages and compared infection to a control strain expressing yellow fluorescent protein (YFP). PUA-Cal-HAD conferred robust protection against multiple phages, including *Tunavirinae* (Bas12), *Tequatrovirus* (T2, Bas36, Bas37, Bas38), and *Tequintavirus* (T5) phages (Figure 2B, **Figure S2A**).

**Figure 2.**
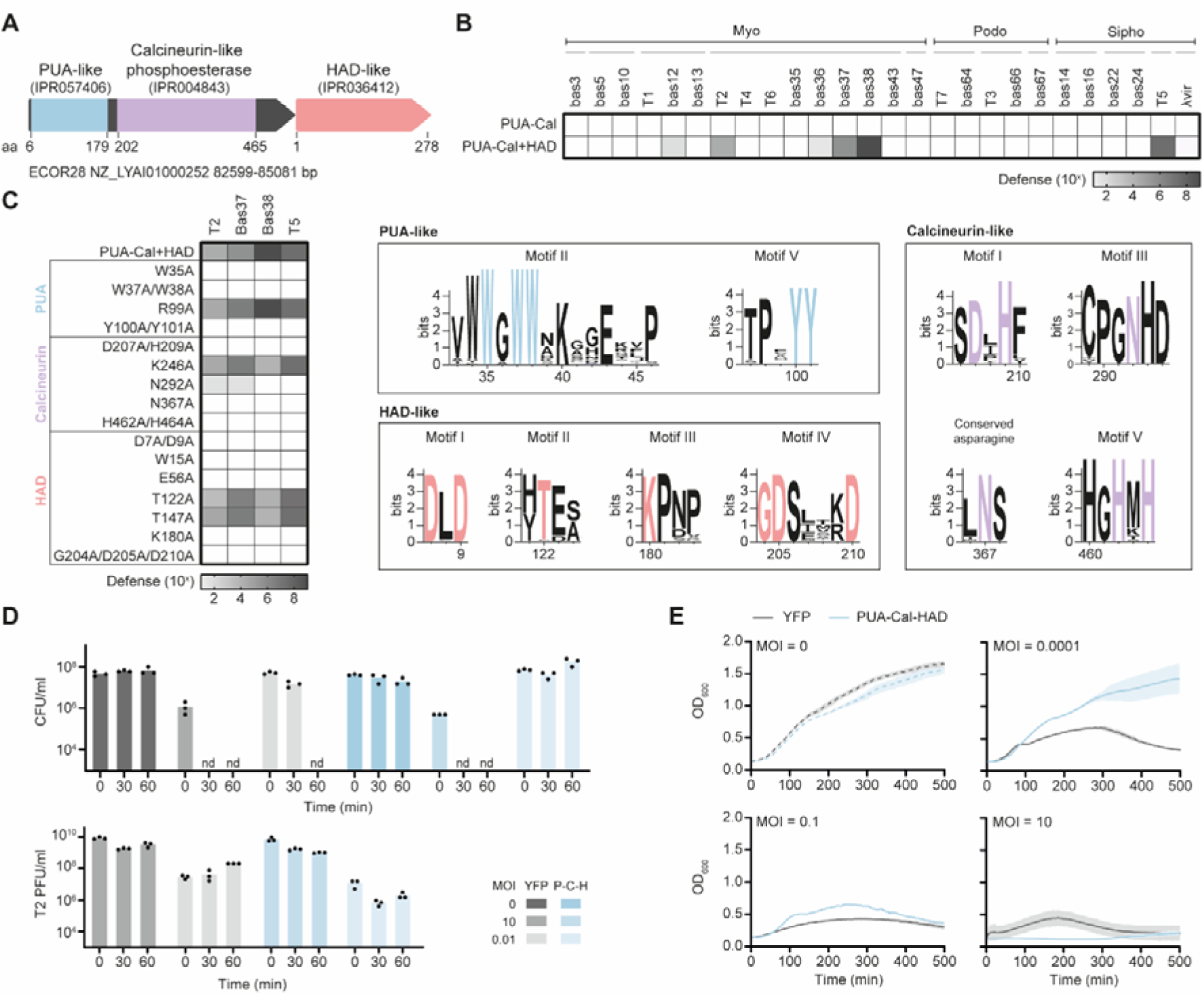
The PUA-Cal-HAD gene cluster provides anti-phage activity. **(A)** The PUA-Cal-HAD system from *E. coli* ECOR28 is composed of two genes with three predicted domains. **(B)** Ten-fold defence of PUA-Cal-HAD against a panel of distinct phages. **(C)** Effect of mutations in conserved motifs in the PUA-like, calcineurin-like phosphoesterase, and HAD-like domains, in defence against phages T2, Bas37, Bas38, and T5. **(D)** Phage (PFU/ml) and bacterial (CFU/ml) concentrations in liquid cultures of control and PUA-Cal-HAD-expressing cells infected with phage T2 at multiplicity of infections (MOI) of 10 and 0.01. **(E)** Growth curves of YFP- and PUA-Cal-HAD-expressing cells infected with phage T2 at different MOIs.

We then tested if all three domains of the ECOR28 locus, namely the N-terminal PUA-like domain (residues 6-179), the fused C-terminal Calcineurin-CE (residues 202-465), and the HAD phosphatase (Figure 2A), are required for the observed defence activity. We introduced mutations in conserved motifs of each domain and assayed protection against T2, Bas37, Bas38, and T5. In the PUA-like domain, a conserved tryptophan-rich motif (motif II, VWWGWWxKxxExxP) and a tyrosine pair in motif V (TPxYY) likely form an aromatic pocket for nucleotide recognition^8^. Consistent with this, substitutions W35A, W37A/W38A, and Y100A/Y101A completely abolished anti-phage activity (Figure 2C**, Figure S2B**). In the Calcineurin-CE domain, we mutated metal-coordinating residues (D207A/H209A, D255A, H420A, and H462A/H464A) and a conserved asparagine (N367A) implicated in substrate recognition^8^. All these mutants lost protection, indicating that catalytic activity of the Calcineurin-CE domain is essential for defence (Figure 2C**, Figure S2B**).

The HAD protein in this system belongs to the type I (C1) HAD phosphatase family, containing the canonical four catalytic motifs and a mobile α-helical cap domain inserted between motifs I and II^23,24^. Guided by the conserved HAD consensus, we mutated the catalytic aspartates in motif I (D7A/D9A)^25–27^, the threonine in motif II (T122A), the lysine in motif III (K180A)^23^, and acidic residues in motif IV (G204A/D205A/D210A)^23,25,28–30^, as well as conserved residues in the cap (W15A, E56A). Each of these mutations (except T122A), and the deletion of the HAD gene entirely, abolished anti-phage defence (Figure 2B**,C****; Figure S2B**). Together, these data show that the PUA-like, Calcineurin-CE, and HAD domains, as well as their ligand-binding and catalytic motifs, are all required for protection.

To understand how this system impacts infection dynamics, we followed T2 infection at a multiplicity of infection (MOI) of 10. In cells expressing PUA-Cal-HAD, both bacterial growth and phage amplification were arrested (Figure 2D**,E**). At a lower MOI of 0.01, PUA-Cal-HAD-expressing cultures survived infection, whereas control cultures were lysed. These MOI-dependent outcomes are consistent with an abortive infection strategy in which activation of the PUA-Cal-HAD system halts phage replication at the cost of host cell viability, thereby protecting the bacterial population.

### PUA-Cal assembles into a metal-stabilised hexameric scaffold

To define the structural organisation of PUA-Cal, we combined AlphaFold 3 (AF3) modelling with extensive cryo-EM and solution biochemistry. AF3 predicts that the PUA-Cal fusion protein adopts a bilobed architecture in which the N-terminal PUA-like domain (1-185) is juxtaposed against the C-terminal Cal phosphoesterase domain (residues 186-549) (Figure 3A, left). Consistent with this prediction, the cryo-EM structure shows that the PUA-like domain adopts a compact α/β fold characteristic of the PUA family^14^ (Figure 3B, top), whereas the Cal domain forms a larger α/β phosphoesterase fold organised around a conserved catalytic core (Figure 3B, bottom). In the AF3 model (Figure 3A, left), the PUA-like domain harbours a deeply recessed aromatic pocket, while the Cal domain likely coordinates its substrate through a conserved binuclear metal-binding centre. These two ligand-binding sites are positioned on opposite faces of the molecule, indicating that ligand recognition by the PUA-like domain is structurally decoupled from catalysis in the Cal domain, while remaining potentially linked through interdomain communication medicated by the flexible PUA-Cal junction.

**Figure 3.**
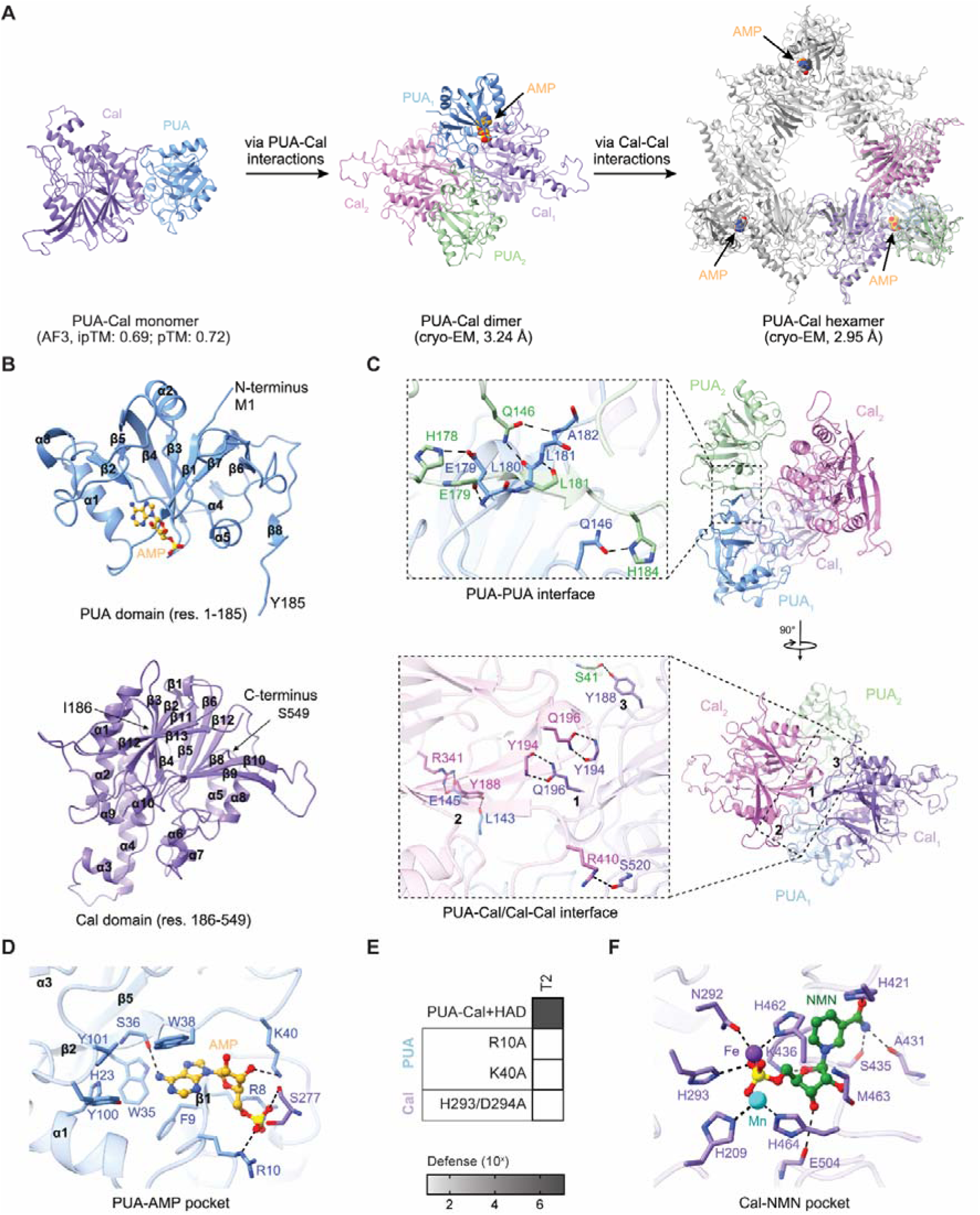
Structural architecture of PUA-Cal. **(A)** Assembly pathway of the PUA-Cal fusion protein. Left, AlphaFold 3 (AF3) model of the PUA-Cal monomer. Middle, cryo-EM structure of the PUA-Cal dimer formed through PUA-Cal interactions. Right, cryo-EM reconstruction of the PUA-Cal hexamer assembled from three dimers via Cal-Cal interfaces, with AMP bound at the PUA-like domains. **(B)** Domain organisation of the PUA-Cal fusion. Top, structure of the PUA-like domain (residues 1-185) with bound AMP. Bottom, structure of the Calcineurin-CE domain (residues 186-549) showing the overall α/β fold. **(C)** Structural details of the interfaces stabilising the PUA-Cal dimer. Top, close-up view of the PUA-PUA interface, highlighting residues that mediate inter-subunit contacts between PUA-like domains. Bottom, close-up view of the composite PUA-Cal/Cal-Cal interface, showing interactions between the PUA-like domain of one subunit and the Cal domain of the neighbouring subunit. Right, overall views of the PUA-Cal dimer in two orientations. **(D)** Close-up view of the AMP-binding pocket in the PUA-like domain, highlighting residues involved in adenine recognition and phosphate coordination. **(E)** Effect of mutations in key residues in the catalytic pockets of the PUA-like and Calcineurin-CE domains on PUA-Cal-HAD defence against phage T2. **(F)** Close-up view of the Calcineurin-CE active site with NMN modelled, showing the dinuclear metal centre and conserved catalytic residues involved in cofactor binding.

Biochemical analysis revealed that PUA-Cal populates multiple oligomeric states in solution, with divalent metal ions strongly stabilising the hexameric assembly. Size-exclusion chromatography showed that, in the absence of added divalent cations, PUA-Cal exists as a mixture of monomeric, dimeric, and hexameric species, whereas addition of cations (Fe^3+^, Mg^2+^, or Mn^2+^) shifts the equilibrium toward a single, well-defined hexameric assembly (**Figure S3A**). These oligomeric assignments are further supported by mass photometry, which independently confirmed the presence of PUA dimers and hexamers in solution **(Figure S3B**).

Cryo-EM reconstruction (**Figure S3C-E**) revealed that the fundamental oligomeric building block of PUA-Cal is an asymmetric dimer stabilised by composite PUA-PUA, PUA-Cal, and Cal-Cal interfaces (Figure 3A, middle; Figure 3C). The PUA-PUA interface is mediated by an extensive network of hydrogen bonds and salt bridges formed by conserved polar and charged residues, centred on a prominent salt bridge between H178 of one protomer and E179 of the opposing protomer, together with Q146, which anchors the two PUA-like domains in a defined relative orientation. Additional inter-subunit hydrogen bonds involving residues within the flexible PUA-Cal junction loop (residues 173-193), including E179 and L181, further stabilise this interface. Adjacent to this, Cal-Cal interactions within the dimer are mediated primarily by mainchain interactions involving residues Y194-K196 within the β1 strand at the N-terminus of the Cal domain from each protomer, together with an additional hydrogen bond formed by R410 and S520 that stabilises the C-terminal region of the Cal-Cal interface.

The PUA-Cal interface is mediated predominantly by loop regions of the Cal domain from each inner protomer. Within the flexible junction loop, Y188 forms a side-chain hydrogen bond with L143 located in the extended loop between α5 and β6 of the PUA-like domain, while E145 forms bifurcated hydrogen bonds with R341 on the Cal domain. The second protomer establishes a related but non-equivalent set of interactions, in which Y188 forms a hydrogen bond with the main-chain atoms of S41 at the β2-α2 boundary of the PUA-like domain, reflecting the intrinsically asymmetric organisation of the dimer interface. Structural superposition of the two protomers based on the Cal domains reveals nearly identical catalytic cores (RMSD ≈ 0.48 Å), whereas the PUA-like domains occupy markedly different spatial positions, with displacements of up to ∼18 Å (**Figure S4A**, left). Conversely, superposition based on the PUA-like domains highlights a corresponding shift of the Cal domains of ∼19 Å (**Figure S4A**, right). This asymmetry arises from flexibility within the interdomain linker while the catalytic cores remain highly conserved.

The PUA-Cal dimers further assemble via Cal-mediated interfaces into a non-planar, ring-like hexamer, the stability of which is enhanced by divalent metal ions (Figure 3A, right). Detailed inspection of the hexameric assembly shows that inter-subunit contacts are dominated by extensive Cal-Cal interfaces structurally preserved across assembly states, which form the rigid structural scaffold of the complex (**Figure S4B**). These interfaces are mediated by complementary β-strand pairings that generate continuous intermolecular β-sheets, reinforced by loop-loop interactions involving loops 372-379, 421-437, and 431-442. Together, these contacts create a contiguous interaction surface that bridges neighbouring Cal domains and stabilises the ring-like architecture, while remaining largely insensitive to conformational changes in the PUA-like domains.

The cryo-EM structural analysis revealed a well-defined AMP-binding pocket within the PUA-like domain (Figure 3D). Clear density corresponding to AMP is observed within a deep aromatic cavity formed by F9, H23, W35, W38, Y100, and Y101, which stabilise the adenine base through π-π stacking and hydrophobic interactions. The ribose and phosphate moieties extend toward the pocket entrance and are anchored by polar interactions, with R8 and R10 forming hydrogen bonds to the phosphate group and K40 engaging the ribose moiety. Accordingly, mutation in these residues (R8A or K40A) abolished defence against phage T2 (Figure 3E). AMP is derived from endogenous cellular pools and is not added during purification or grid preparation, indicating that AMP binding reflects a physiologically relevant basal state. Comparison across monomeric, dimeric, and hexameric assemblies shows that the geometry of the AMP-binding pocket is conserved, suggesting that AMP binding precedes and is independent of higher-order oligomerisation.

Reconstructions of AMP-bound PUA-Cal hexamers revealed two predominant conformation states (**Figure S4C**). These states correlate with partial occupancy of the PUA nucleotide-binding pockets, as only three AMP molecules are bound per hexamer, with each dimer contributing a single AMP-bound PUA-like domain. In the ‘2-up, 1-down’ state, two AMP molecules are bound to inward-facing PUA-like domains, whereas the third is bound to an outward-facing PUA-like domain. In the ‘3-down’ state, all three AMP molecules are bound to inward-facing PUA-like domains. Superposition of these states shows that the overall hexameric architecture is preserved (RMSD ≈ 0.85 Å), with conformational variability localised almost exclusively to the PUA-like domains and the flexible junction loop spanning residues 173-193 (**Figure S4D,E**), consistent with an autoinhibited basal state.

Structural analysis of ion-bound PUA-Cal hexamers revealed an architecture indistinguishable from the AMP-bound state (**Figure S4F,G**), indicating that metal ions stabilise a pre-existing assembly rather than redefining hexamer organisation. Ion binding induces subtle tightening of Cal-Cal contacts and local rearrangements in surface-exposed loops without perturbing the geometry of the catalytic metal centre of the PUA-Cal junction (**Figure S4H,I**).

Across all basal states, the Cal domain maintains a conserved catalytic architecture that closely resembles canonical metallo-dependent phosphoesterases, centred on a stably nicotinamide mononucleotide (NMN)-occupied pocket (Figure 3F**, Figure S5A**). NMN is coordinated by a binuclear metal centre formed by conserved histidine residues (H209, H293, H462, and H464), with the phosphate moiety oriented toward the metal ions in a catalytically competent configuration. Structural modelling further indicates that the Cal catalytic pocket can accommodate other adenine-containing nucleotides, including NAD^+^ and ATP, which bind in similar orientations with their phosphate groups positioned adjacent to the metal centre (**Figure S5B,C**), indicating that the Cal domain is structurally capable of binding multiple nucleotide substrates.

Collectively, these data define the PUA-Cal hexamer as a metal-stabilised, conformationally plastic basal state in which a rigid Cal catalytic scaffold is coupled to a flexible, ligand-responsive PUA module, positioning the complex for ligand-dependent activation.

### The HAD protein associates with the PUA-Cal hexamer

Our cryo-EM data show that PUA-Cal assembles into a cation-stabilised hexamer with a central Calcineurin-CE core (Figure 3A). Because the HAD protein is essential for defence and its domain architecture suggests a potent phosphatase, we next asked whether it associates with this hexameric scaffold.

Biochemical analyses revealed that the HAD phosphatase behaves as a strictly monomeric protein in solution (**Figure S5D,E**). Despite several attempts to obtain a structure of the protein, HAD consistently aggregated preventing structural analysis. Using AF3, we observed that HAD is a structurally autonomous enzyme adopting the canonical HAD phosphatase topology, with a central Rossmannoid α/β core flanked by auxiliary helical elements (Figure 4A, left). The catalytic core is stabilised by an extensive network of hydrogen bonds and salt bridges and forms a well-defined active-site pocket shaped by conserved polar residues and an aromatic cage that positions the substrate for catalysis. Central to this architecture is the hallmark DxDx(T/N) catalytic motif, in which the invariant D7 and D9 residues (essential for defence, Figure 2C) coordinate a catalytic Mg^2+^ ion and constitute the nucleophilic centre required for phosphotransfer (Figure 4A, middle and right).

**Figure 4.**
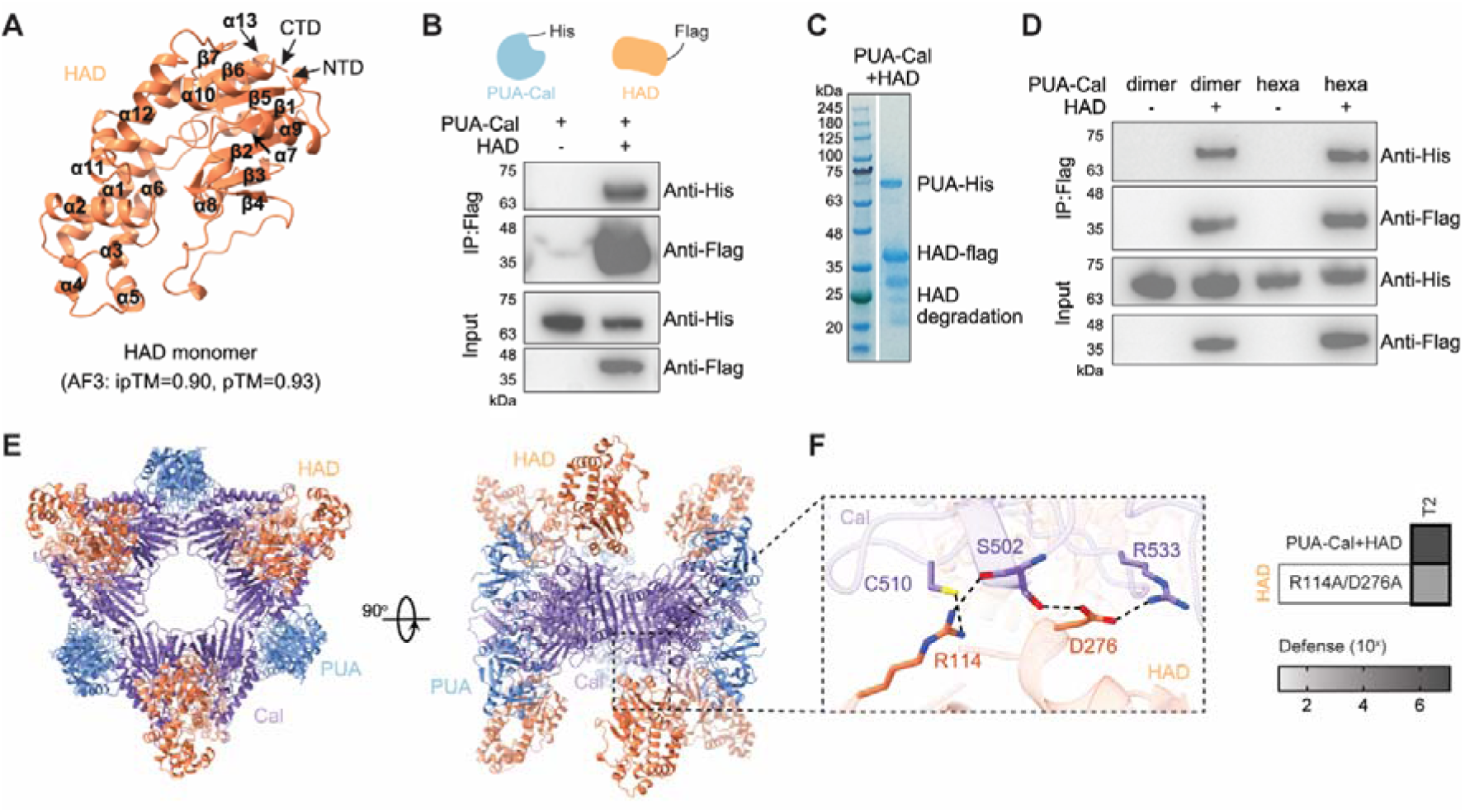
HAD is a catalytically competent phosphatase that directly associates with the PUA-Cal complex. **(A)** AlphaFold 3 (AF3) model of the HAD protein bound to AMP and Mg2+, showing a canonical haloacid dehalogenase fold. Inset highlights the conserved DxDx(T/N) catalytic motif. **(B)** Co-immunoprecipitation assay with HAD-Flag and PUA-Cal-His. **(C)** SDS-PAGE analysis of the co-IP shown in (B), confirming recovery of both PUA-Cal-His and HAD-Flag in the immunoprecipitated fraction. Partial degradation of HAD is indicated. **(D)** In vitro pull-down assays testing direct interaction between HAD and dimeric and hexameric PUA-Cal assemblies. **(E)** AF3 model of the PUA-Cal-HAD complex arranged as a hexamer, shown in two orientations. HAD molecules are positioned on the outer surface of the PUA-Cal assembly and contact the Cal domains. **(F)** Close-up view of the predicted Cal-Had interface. HAD is positioned distal to the Cal active site. Right, effect of mutations in HAD residues involved in the interaction with PUA-Cal, in the PUA-Cal-HAD activity against phage T2 infection.

Structural superposition reveals that HAD closely resembles the catalytic core of human Pdxp/Chronophin, with an overall RMSD of approximately 0.93 Å, indicating strong conservation of both fold and active site geometry (**Figure S5F-H**). In both enzymes, the Rossmannoid fold positions the DxDx(T/N) motif at the base of the active site, where it coordinates Mg^2+^ and stabilises the transition state during catalysis, supporting classification of HAD as a bona fide member of the HAD phosphatase family.

Co-immunoprecipitation, pull-down assays, and mass photometry demonstrate that HAD associates directly with PUA-Cal dimers and hexamers, although the interaction is relatively weak and the resulting complexes are prone to aggregation in solution, precluding high-resolution structural determination by cryo-EM (Figure 4B-D**, Figure S3B, Figure S5I,J**). To gain insight into the architecture of the PUA-Cal-HAD complex, we modelled the full assembly using AlphaFold3. The model positions six HAD molecules around the periphery of the PUA-Cal hexamer, contacting Cal domains at dimer-dimer interfaces (Figure 4E). The interaction is mediated primarily by polar contacts and salt bridges, including a representative salt bridge between HAD residue D276 and Cal residue R533, together with additional contacts involving Cal residue S502. Further stabilisation is provided by hydrogen bonds between Cal residues C510 and S502 and HAD residue R133, collectively defining a weak but specific interaction interface (Figure 4F). Accordingly, mutation R114A/D276A in HAD significantly reduced defence against phage T2 (Figure 4F). In this configuration, HAD active sites face outward and are separated from the Cal catalytic centres by approximately 36 Å, arguing against direct substrate channelling. The HAD-Cal interface is spatially distinct from the Cal catalytic pocket and does not occlude the active site, indicating an association that preserves Cal catalytic integrity and is consistent with a sequential rather than channelled two-enzyme mechanism.

Together, these observations suggest that HAD interacts with the PUA-Cal hexamer and that this interaction is required for anti-phage defence.

### PUA-Calcineurin-HAD depletes the cellular deoxyribonucleotide triphosphate pool

The effector activity of the PUA-Cal-HAD defence system is thought to reside in its calcineurin-CE and HAD domains, which are predicted to degrade cellular nucleotide-based metabolites^8^. Unlike classical calcineurins that primarily dephosphorylate protein substrates, the calcineurin-CE family is more distantly related and has been proposed to act on nucleotide substrates, such as mono- and dinucleotides, NAD^+^, cyclic nucleotides, and/or polynucleotides^8^. Phylogenetic analyses support a functional divergence toward (oligo)nucleotide substrates^8^. Together, these observations suggested that PUA-Cal-HAD might protect cells by depleting nucleotide or nucleotide-derived pools during infection.

To test this, we quantified intracellular nucleotide and cofactor levels by LC-MS in BL21-AI strains expressing YFP (control) or PUA-Cal-HAD, following infection with T2. We measured NTPs, dNTPs, NAD^+^ and NMN at 15 minutes post-infection. Cells expressing the PUA-Cal-HAD system displayed a 4-20-fold depletion of dNTPs compared to the YFP control cells (Figure 5A**, Figure S6A, Table S2**). These findings support a model in which, once activated, the PUA-Cal-HAD system rapidly perturbs cellular deoxyribonucleotide pools, enforcing a state of nucleotide starvation that prevents productive phage replication and arrests host growth.

**Figure 5.**
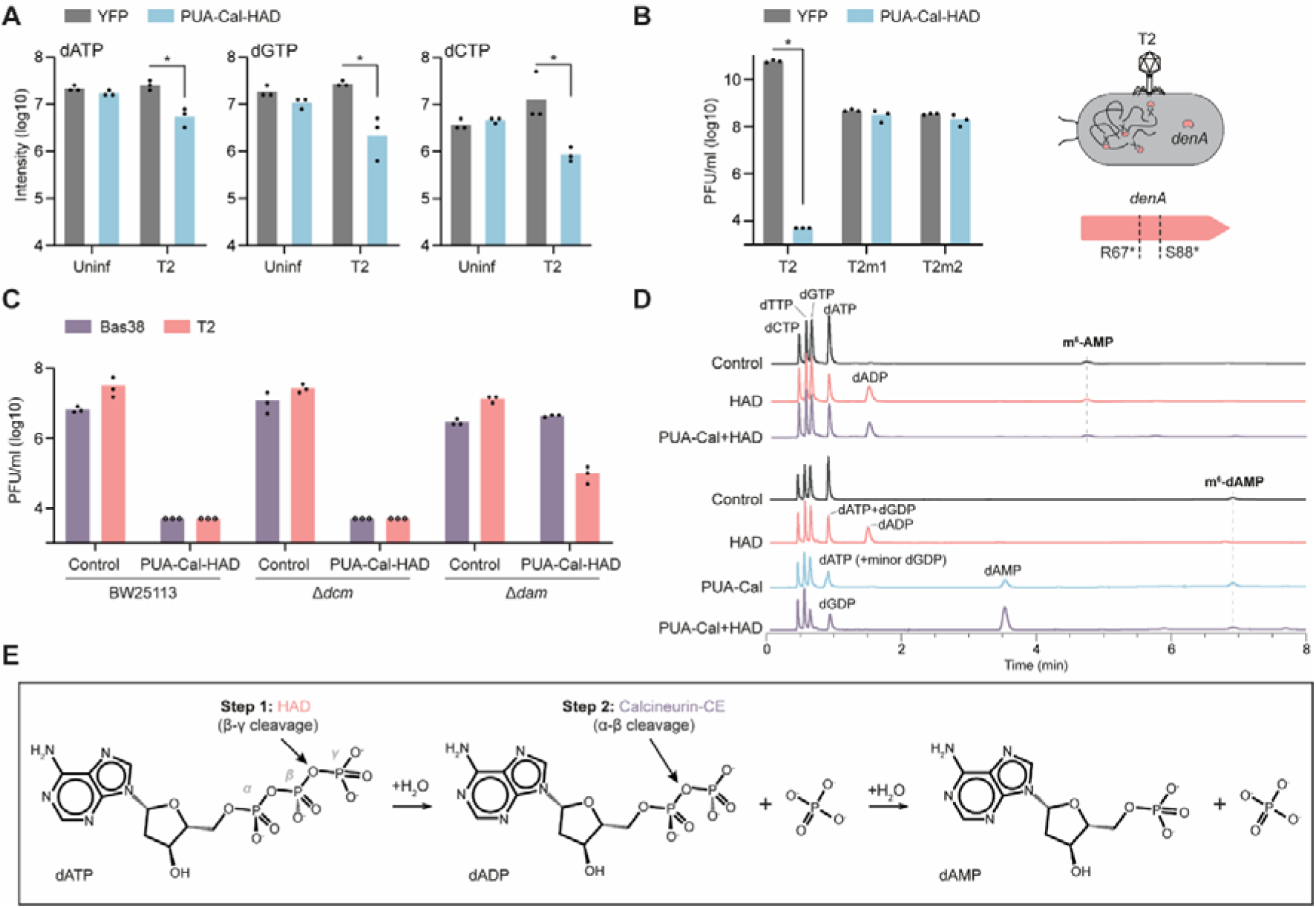
PUA senses m^6^-dAMP to activate nucleotide depletion by Cal and HAD. **(A)** Quantification of nucleotide metabolites by LC-MS, shown as relative intensity levels in log10, in cells expressing YFP or PUA-Cal-HAD upon infection with phage T2. **(B)** Infectivity of T2 phage escape mutants on YFP- and PUA-Cal-HAD-expressing cells. Escape mutants have mutations in gene *denA*. **(C)** Infectivity of Bas38 and T2 phages on YFP (control) or PUA-Cal-HAD cells containing both methylases Dcm and Dam (BW25113), or versions with deleted methylases. **(D)** LC-MS measurement of PUA-Cal-HAD *in vitro* activity on dNTPs in the presence of m^6^-AMP or m^6^-dAMP. **(E)** Reaction scheme for PUA-Cal-HAD activity. Step 1: HAD hydrolyses the β-γ phosphoanhydride bond of dATP to generate dADP and inorganic phosphate (Pi). Calcineurin-CE then hydrolyses the α-β phosphoanhydride bond of dADP to yield dAMP and Pi.

### PUA senses methylated dAMP generated during phage-induced host DNA degradation

To determine what triggers activation of the PUA-Cal-HAD system during infection, we isolated T2 escape mutants that retained infectivity in PUA-Cal-HAD-expressing cells. Whole-genome sequencing revealed that all escapers carried frameshift mutations in *denA*, encoding the T4-family endonuclease responsible for degrading host DNA early during infection (EndoII) (Figure 5B**, Table S3**). EndoII cleaves host DNA into fragments that are further processed by Endo IV (*denB*) into deoxyribonucleoside monophosphates (dNMPs), which are recycled into phage DNA synthesis^31^. Although DenA is not strictly essential for T2 replication, it enhances replication efficiency by increasing nucleotide availability^32^. The enrichment of *denA* mutations in escape phages therefore indicates that phage-mediated degradation of host DNA is required for PUA-Cal-HAD activation.

The N-terminal domain of PUA-Cal belongs to the PUA/ASCH clan, whose members commonly recognise nucleotides or modified nucleobases via conserved nucleobase-binding pockets enriches in aromatic residues, rather than relying solely on backbone contacts^33,34^. This domain architecture led us to hypothesise that PUA-Cal-HAD might not simply sense DNA breaks^35^ but instead respond to a modified nucleotide generated when the host chromosome is degraded. In *E. coli*, the predominant chromosomal modifications are N^6^-methyladenine (m^6^dA), installed by the Dam methyltransferase, and 5-methylcytosine (m^5^dC), installed by Dcm. We therefore asked whether either of these methylation systems is required for PUA-Cal-HAD-mediated defence.

To test this, we expressed PUA-Cal-HAD under its native promoters in strains from the KEIO collection, including wild-type BW25113 (*dam*⁺ *dcm*⁺), JW3350 (*dam*⁻ *dcm*⁺) and JW1944 (*dam*⁺ *dcm*⁻) and assayed anti-phage activity. The system remained active in wild-type and *dcm*⁻ backgrounds but showed abolished protection against Bas38 and markedly reduced protection against T2 in *dam*⁻ cells (Figure 5C). T2, but not Bas38, encodes an N^6^-adenine methyltransferase (Genbank accession no. XUQ82749.1), which may account for the residual protection observed in a *dam^-^* background. Thus, activation of PUA-Cal-HAD depends on Dam-mediated N^6^-adenine methylation, but not on Dcm-mediated cytosine methylation.

The combined DenA and Dam dependencies suggested two possible models for activation: PUA-Cal-HAD could sense (i) damaged, Dam-methylated DNA fragments, or (ii) a modified nucleotide derived from Dam-methylated DNA. To test the first possibility, we examined DNA binding by PUA-Cal to linear substrates carrying different methylation patterns. Electrophoretic mobility shift assays showed that PUA-Cal binds linear single-stranded and double-stranded DNA efficiently, but does so promiscuously, with no detectable discrimination between unmethylated DNA and DNA containing m^6^A or m^5^C (**Figure S6B**). Thus, although PUA-Cal can bind DNA, specific recognition of methylated DNA ends is unlikely to be the primary trigger for activation.

The PUA-Cal structure provided an alternative clue. In the non-infected state, the PUA pocket contains clear density for an AMP molecule bound in the aromatic cavity, and mutations in residues that contact this ligand (R10A, K40A) abolished defence (Figure 3E). Because the protein was purified from uninfected cells, in which DNA degradation is minimal and free modified NMPs and dNMPs are expected to be scarce, and because we established the dependency on Dam methylation, we reasoned that AMP likely represents a basal ligand occupying the pocket in the off state, while a methylated adenine nucleotide produced during infection could be the true activating substrate. When Dam-methylated DNA is degraded to dNMPs during T2 infection, it generates N^6^-methyl-deoxyadenosine monophosphate (m^6^-dAMP), suggesting this nucleotide as a strong candidate for the activating signal.

Because deoxyribonucleotide depletion provides a robust readout of PUA-Cal-HAD activation, we used this as a functional assay to test whether purified PUA-Cal-HAD can be directly activated by candidate adenine nucleotides *in vitro*. We incubated PUA-Cal-HAD with dNTPs and added either m^6^-AMP or m^6^-dAMP as potential ligands, then quantified nucleotide levels by LC-MS. We observed that m^6^-dAMP induced dNTP depletion, whereas m^6^-AMP had no effect (Figure 5D).

Altogether, these results show that PUA senses m^6^-dAMP derived from phage-mediated host genome degradation to activate the PUA-Cal-HAD anti-phage response.

### PUA-Cal-HAD is a coupled effector module in which HAD converts dNTPs into dNDPs and Calcineurin-CE completes conversion to dNMPs

Our metabolomics analysis in infected cells showed that activation of PUA-Cal-HAD leads to a rapid collapse of dNTP pools (Figure 5A). This phenotype indicates that the effector output of the system is enzymatic depletion of dNTP substrates. We therefore asked whether the enzymatic activities of the Calcineurin-CE and HAD components, measured *in vitro*, could account for the selective metabolite changes observed *in vivo*.

For this, we performed LC-MS analysis of reactions containing purified PUA-Cal, with or without HAD, and defined nucleotide substrates in the presence of the activating ligand m^6^-dAMP. HAD alone displayed robust activity on dATP and dGTP, converting them to dADP and dGDP, respectively (Figure 5D), consistent with a dNTP γ-phosphatase activity that hydrolyses the terminal γ-phosphate (Figure 5E). In contrast, PUA-Cal alone showed little to no detectable turnover of dNTPs under these conditions (Figure 5D). Importantly, when PUA-Cal and HAD were combined, the reactions proceeded to completion, with conversion of dATP into dAMP (Figure 5D). Consistent with a stepwise pathway, dADP accumulated when HAD was present alone, whereas these intermediates were depleted in the presence of PUA-Cal, indicating that Calcineurin-CE consumes the dNDP products generated by HAD to produce dNMPs. This activity is consistent with a dNDP phosphohydrolase that hydrolases the α-β phosphoanhydride bond, removing the terminal β-phosphate of the diphosphate (Figure 5E).

We next tested additional substrates to define the effector range of the complex. With ribonucleotides, we observed the same behaviour, with HAD converting ATP to ADP, while efficient accumulation of AMP required the presence of PUA-Cal (**Figure S6C**). In contrast, PUA-Cal catalysed robust cleavage of NAD^+^ to NMN and AMP, and this activity was completely independent of HAD (**Figure S6C**). Moreover, HAD did not reduce or otherwise alter NAD turnover by PUA-Cal, indicating that HAD does not act as a general inhibitor or substrate-gating factor for the Calcineurin-CE active site *in vitro*.

These substrate profiles provide a mechanistic explanation for the two-protein architecture. For dNTP (and ATP) substrates, our data support a sequential dephosphorylation cascade in which HAD first hydrolases the β-γ bond to generate (d)NDP + Pi, and Calcineurin-CE then hydrolyses the α-β bond to generate (d)NMP + Pi (Figure 5E). By contrast, for NAD^+^, Calcineurin-CE cleaves NAD^+^ into two mononucleotide products (NMN and AMP) via hydrolysis of its internal pyrophosphate linkage, and this reaction proceeds without any requirement for HAD.

AF3 modelling was used to further examine how nucleotide binding and HAD engagement are structurally accommodated within the PUA-Cal-HAD system and how these interactions related to the assembly states observed by cryo-EM. Because AF3 currently does not support modelling with dNTP ligands, we used the corresponding ribonucleotide analogues (ATP for HAD and ADP for Cal) as proxies for the structural analyses below. AF3 predicts that HAD adopts a canonical HAD phosphatase fold and binds ATP-Mg^2+^ within a conserved catalytic pocket centred on the DxDx(T/N) motif, in which the invariant D7 residue coordinates the catalytic metal ion and D9 engages the phosphate group, positioning the nucleotide in a catalytically competent configuration (**Figure S6D**). In parallel, AF3 models of the PUA-Cal complex show that ADP and AMP can be simultaneously accommodated within the Cal and PUA binding pockets, respectively, without steric conflict and with relative domain orientations closely resembling those observed in basal cryo-EM assemblies (**Figure S6E**). In this configuration, ADP is positioned adjacent to the Cal binuclear metal centre, consistent with its potential role as a Cal substrate, while AMP occupies the conserved aromatic pocket of the PUA domain. Together, these models indicate that upstream nucleotide processing by HAD and nucleotide engagement by Cal are structurally compatible in the complex.

Together with the *in vivo* metabolomics data, these results support a two-enzyme effector cascade in which activation of PUA-Cal-HAD by Dam-derived m^6^-dAMP unleashes coordinated destruction of dNTP pools. HAD provides the initiating dNTP to NDP activity, while Calcineurin-CE completes nucleotide depletion by converting dNDPs to dNMPs. This coupled mechanism provides a mechanistic basis for the rapid collapse of cellular dNTP pools during infection and, consequently, for the abortive infection phenotype imposed by the PUA-Cal-HAD system.

### m^6^-dAMP converts PUA-Cal from an autoinhibited hexamer into a filamentous effector

To determine the conformational changes occurring upon activation of the defence system by sensing of m^6^-dAMP by the PUA-like domain, we added the ligand to purified PUA-Cal and examined the assemblies formed by cryo-EM. Addition of m^6^-dAMP induced a pronounced reorganisation of PUA-Cal into extended filamentous structures (Figure 6A**,B**, **Figure S7A,B**). These filaments were not detected in the presence of m^6^-AMP or m^6^A-containing single-stranded DNA (**Figure S7A,B**), indicating that filament formation is specifically triggered by m^6^-dAMP and not a generic consequence of adenine nucleotide binding, and correlates with effector activation.

**Figure 6.**
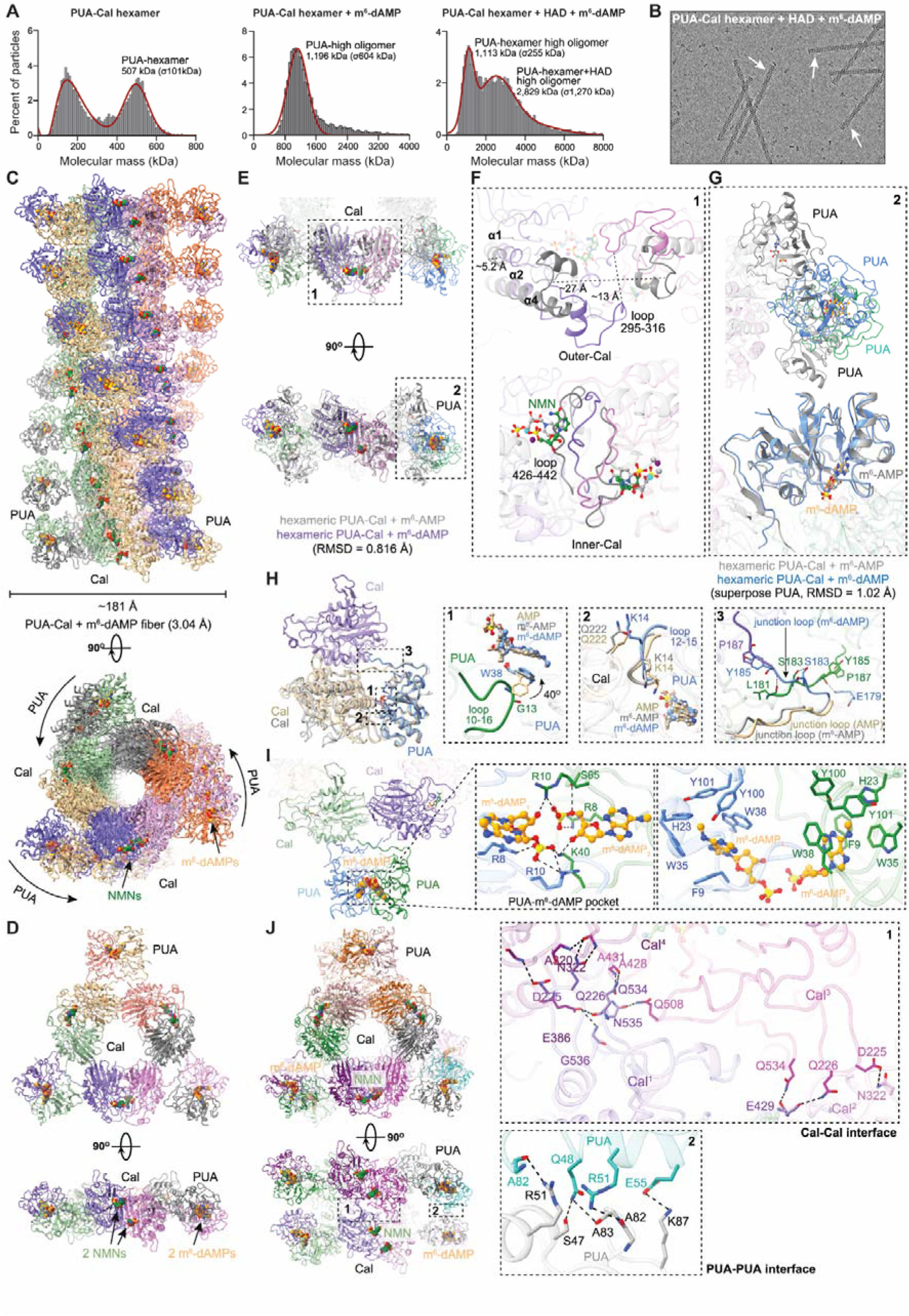
Sensing of m^6^-dAMP by PUA drives PUA-Cal hexamer reorganisation and filament formation. **(A)** Molecular mass distributions measured by mass photometry for purified PUA-Cal in the absence and presence of HAD and m^6^-dAMP. Data are shown as the percentage of particles per molecular mass (kDa). **(B)** Representative cryo-EM micrograph of PUA-Cal mixed with HAD and m^6^-dAMP, showing filamentous assemblies (white arrows). **(C)** Cryo-EM reconstruction of the m^6^-dAMP bound PUA-Cal filament. Top, side view showing stacked PUA-Cal hexamers forming an extended fibre with an axial repeat of ∼181 Å. Bottom, top view illustrating PUA-like domains on the outer surface and the Cal domains forming the inner core. **(D)** Structure of a single m^6^-dAMP bound PUA-Cal hexamer extracted from the filament, shown in two orientations. m^6^-dAMP is bound in each PUA-like domain (six ligands per hexamer), while NMN occupies the Cal active sites. **(E)** Superposition of m^6^-dAMP and m^6^-AMP bound PUA-Cal hexamers, highlighting ligand-dependent conformational differences. Close up views of Cal and PUA-like domains are shown in (F) and (G), respectively. **(F)** Close-up view of the Cal dimer comparing m^6^-dAMP and m^6^-AMP bound states. In the m^6^-dAMP bound state, the two Cal subunits within each dimer move closer together (inward displacements of helices α1, α2, and α4 of up to ∼5.2 Å). The loop spanning residues 295-316 shifts toward the dimer interface, reducing the distance between opposing loops from ∼27 Å to ∼13 Å. This arrangement is accompanied by a reorientation of bound NMN driven by movement of loop 426-442, while coordination of the catalytic metal centre and the overall active-site geometry remain conserved. **(G)** Superposition of PUA-like domains from m^6^-dAMP and m^6^-AMP bound hexamers. In the m^6^-dAMP-bound state, PUA-like domains undergo pronounced rotations/translation shifts, bringing the two PUA protomers within each dimer into closer apposition and accommodating paired ligand binding. **(H)** m^6^-dAMP-dependent conformational rearrangements at the PUA-Cal interface. Superpositions highlight ligand-dependent interdomain changes: (1) ligand-induced rearrangement of W38 in the PUA nucleotide-binding pocket; (2) m^6^-dAMP specific remodelling of the PUA N-terminal loop (residues 10-16), including reorientation of K14; (3) Ligand-dependent remodelling of the PUA-Cal junction loop (residues 173-193), consistent with an interdomain relay that stabilises an assembly-competent configuration. **(I)** Structural basis of m^6^-dAMP recognition by the PUA-like domain. Top: overall structure of a PUA-Cal dimer within the filament, highlighting m^6^-dAMP bound within each PUA-like domain. Middle: close-up view showing two m^6^-dAMP molecules bound symmetrically within the PUA dimer and coordinated by a Mg^2+^ ion, with phosphate groups stabilised by basic residues (including R10 and K40). Bottom: detailed view of the PUA-like binding pocket, in which the adenine bases are stabilised by π–π stacking with conserved aromatic residues (F9, H23, W35, W38, Y100, and Y101). **(J)** Overall architecture of staked PUA-Cal hexamers forming a continuous filament, shown in two orientations. Insets highlight filament-stabilising interfaces: (1) Cal-Cal contacts between adjacent hexamers formed by hydrogen-bonding and salt bridge networks; (2) PUA-PUA interactions between neighbouring hexamers, showing polar contacts between PUA protomers that are engaged in the filament state but not observed in basal hexameric assemblies.

Cryo-EM analysis revealed long, regular filaments composed of stacked PUA-Cal hexamers, indicating that m^6^-dAMP triggers a transition from a discrete hexameric state to a higher-order polymeric assembly (Figure 6C**,D**). Helical reconstruction of the filament yielded a 3.04 Å map (Figure 6C, **Figure S7C-E**) revealing that the filament is organized as a repeating stack of hexameric rings built from PUA-Cal dimers as the asymmetric unit (**Figure S7F**). Cryo-EM reconstructions of the filament state show that individual hexamers retain their internal Cal-Cal scaffold and overall ring-like architecture but stack through defined inter-hexamer interfaces (Figure 6E-G). Unlike the basal hexamer, in which PUA-like domains adopt mixed ‘up’ and ‘down’ conformations, m^6^-dAMP-bound filaments display a strong bias toward a uniform PUA configuration that forms a coplanar interaction surface, thereby promoting head-to-tail stacking of adjacent hexamers (Figure 6D**,E**). Consistent with this, filament formation correlates with constrained conformations of the PUA-Cal junction loop (residues 173-193), suggesting that ligand binding reduces junction-loop flexibility and stabilises a filament-competent interdomain geometry (Figure 6H, **Figure S7F**). By contrast, the basal hexamer is unable to form filaments because heterogeneous PUA orientations introduce steric clashes that prevent productive inter-hexamer association (**Figure S7G**).

To understand why m6-dAMP specifically promotes filamentation, we compared local and interdomain conformational responses to m^6^-dAMP versus m^6^-AMP and AMP (Figure 6H). Engagement of either m^6^-AMP or m^6^-dAMP induces a shared pocket-level rearrangement centred on W38: ligand binding promotes an ∼40° rotation of the W38 side chain, repositioning the indole ring to stack more parallel to the adenine base and constricting the aromatic pocket relative to the AMP-bound basal state (Figure 6H, panel 1). This reconfiguration optimises π-π stacking and relieves a steric conflict with the opposing PUA N-terminal loop (residues 10-16), enabling closer apposition of two PUA domains. However, only m^6^-dAMP triggers and additional conformational response in the N-terminal loop of the same protomer (Figure 6H, panel 2). In the m^6^-dAMP-bound state, residues L12-S15 shift away from the nucleotide and Lys14 reorients outward, creating a local steric incompatibility Q222 of the adjacent Cal domain. This interdomain conflict is resolved by remodelling of the PUA-Cal junction loop, which reorients the relative PUA-Cal geometry to stabilise a paired, m^6^-dAMP-bound PUA dimer configuration competent for higher-order assembly (Figure 6H, panel 3). Collectively, these data indicate that m^6^-dAMP couples local nucleotide recognition to a junction-loop relay that remodels the PUA-Cal interface, thereby converting ligand binding into a filament-competent architecture.

Mass photometry and solution assays further support this model, showing a shift from discrete oligomeric species to high molecular weight assemblies upon addition of m^6^-dAMP (**Figure S7A**). Filament formation occurs rapidly and is accompanied by visible precipitation at higher protein concentrations, consistent with strong inter-hexamer interactions and cooperative assembly. Structural analysis of the filament state reveals that six m^6^-dAMP molecules are bound per PUA-Cal hexamer (Figure 6D), in contrast to the basal hexamer, in which only three AMP molecules are observed (Figure 3A). Within each PUA dimer, two m^6^-dAMP molecules are positioned in close proximity, facing one another across the dimer interface and coordinated by a Mg^2+^ ion, forming a symmetric nucleotide-metal-nucleotide arrangement (Figure 6I). In this configuration, R10 and K40 from a single PUA protomer contact both ligands, providing a structural basis for cooperative nucleotide sensing and full pocket occupancy (Figure 6I). By comparison, the incomplete and asymmetric AMP occupancy observed in basal hexamers likely reflects an autoinhibited state in which PUA-like domains remain conformationally heterogeneous and unable to engage in productive inter-hexamer contacts.

Importantly, filament assembly does not disrupt the integrity of the Cal catalytic core, and the Cal active site remains occupied by NMN in both the basal hexamer and the m^6^-dAMP-induced filament state. We interpret NMN as a non-physiological, co-purifying ligand that reports on active-site architecture rather than on engagement with the physiological substrates (dNTPs). In both basal and filament states, NMN is coordinated by the conserved binuclear Fe/Mn centre of the Cal domain, with its phosphate group oriented towards the metal ions in a catalytically competent configuration (**Figure S7F**, panel 2). Cal-Cal interfaces within each hexamer remain largely invariant across basal and filament states (**Figure S7F**, panel 3), and no large-scale rearrangements are observed in the catalytic metal centre or NMN-occupied active-site pocket. Filament formation instead tightens local inter-subunit packing. In the filament state, the NMN base undergoes an ∼46° rotation and becomes additionally stabilised by π–π stacking with W316 from a neighbouring Cal subunit within the same hexamer (**Figure S7F**, panel 2), likely reinforcing ligand retention while preserving active site geometry. At the level of dimer organization, the two Cal subunits within each dimer move closer together in the m^6^-dAMP-bound state, primarily through inward displacements of helices α1, α2, and α4 of up to ∼5.2 Å, resulting in tighter inter-subunit packing (Figure 6F). Concomitantly, the loop spanning residues 295-316 shifts towards the dimer interface, reducing the distance between opposing loops from ∼27 Å to ∼13 Å. This compaction is accompanied by a reorientation of the bound NMN molecule, driven by movement of loop 426-442, which rotates predominantly around the phosphate group while preserving metal coordination and overall active site geometry. Higher-order assembly is stabilised by ligand-dependent remodelling of interface surfaces rather than by changes to the catalytic site itself. This filament formation is built upon a rigid Cal scaffold, with polymerisation driven by ligand-dependent reconfiguration of the regulatory PUA-like module and by the emergence of new intra- and inter-hexamer contacts (Figure 6J**, Figure S7F**). Within each hexamer, m^6^-dAMP binding promotes a newly engaged PUA-PUA interface, mediated by helix α2 of one protomer and the extended β4-β5 loop of the adjacent protomer (Figure 6J**, Figure S7F**). Adjacent hexamers are then linked through Cal-Cal interfaces comprising extensive hydrogen-bonding and salt-bridge networks between loops and helices on opposing Cal domains (Figure 6J), providing a structural basis for directional hexamer stacking and filament growth.

Finally, mass photometry analysis indicates that HAD remains associated with PUA-Cal following m^6^-dAMP activation and filament formation (**Figure S7A**). However, superposition of the AF3-predicted PUA-Cal-HAD assembly (**Figure S7H**) and the AF3 model incorporating ATP/ADP/AMP and ions as ligand proxies (**Figure S7I**) onto the m^6^-dAMP-induced filament reveals extensive steric clashes between HAD and neighbouring Cal domains, indicating that HAD cannot retain its basal positioning and must substantially reposition within the activated filament state. These clashes are consistent with a model in which HAD is recruited to a basal PUA-Cal scaffold but undergoes spatial reorganisation upon polymerisation to remain engaged with the complex while enabling access to its nucleotide substrates.

Together, these data indicate that m^6^-dAMP acts as a specific activating ligand that converts the PUA-Cal complex from an autoinhibited, partially occupied basal hexamer into a fully occupied, Mg^2+^-coupled filamentous assembly. In this model, m^6^-dAMP binding elicits a junction-loop-mediated interdomain relay that biases the regulatory PUA-like domains into a uniform, assembly-competent configuration, enabling cooperative filament growth without remodelling the Cal catalytic core. The data further suggest that HAD is recruited to an inactive PUA-Cal scaffold but undergoes spatial reorganisation upon activation, likely allowing it to execute the downstream effector reaction together with Cal.

### Phage DNA mimics antagonise PUA-Cal-HAD by disrupting PUA-mediated assembly

Given the broad DNA-binding capacity of PUA-Cal (**Figure S6B**), we asked whether DNA mimic proteins could antagonise PUA-Cal-HAD. We tested a panel of previously characterised DNA mimic proteins encoded by mobile genetic elements, including Arn^36^, Gam^37^, Ocr^38^, AcrIF2^39,40^, AcrIF18^40,41^, Ugi^42^, P56^43^, Gp44^44^, and ArdA^45^, which represent structurally and mechanistically diverse antagonists of host nucleases and DNA-binding complexes. In plaque assays against phage T5, several of these proteins, including Arn, Gam, Ocr, Ugi, and AcrIF2, reduced or abolished PUA-Cal-HAD-mediated protection (Figure 7A). Using Gam expressed from Anderson promoters at different strengths, we observed dose-dependent inhibition, consistent with a direct antagonistic interaction (Figure 7B).

**Figure 7.**
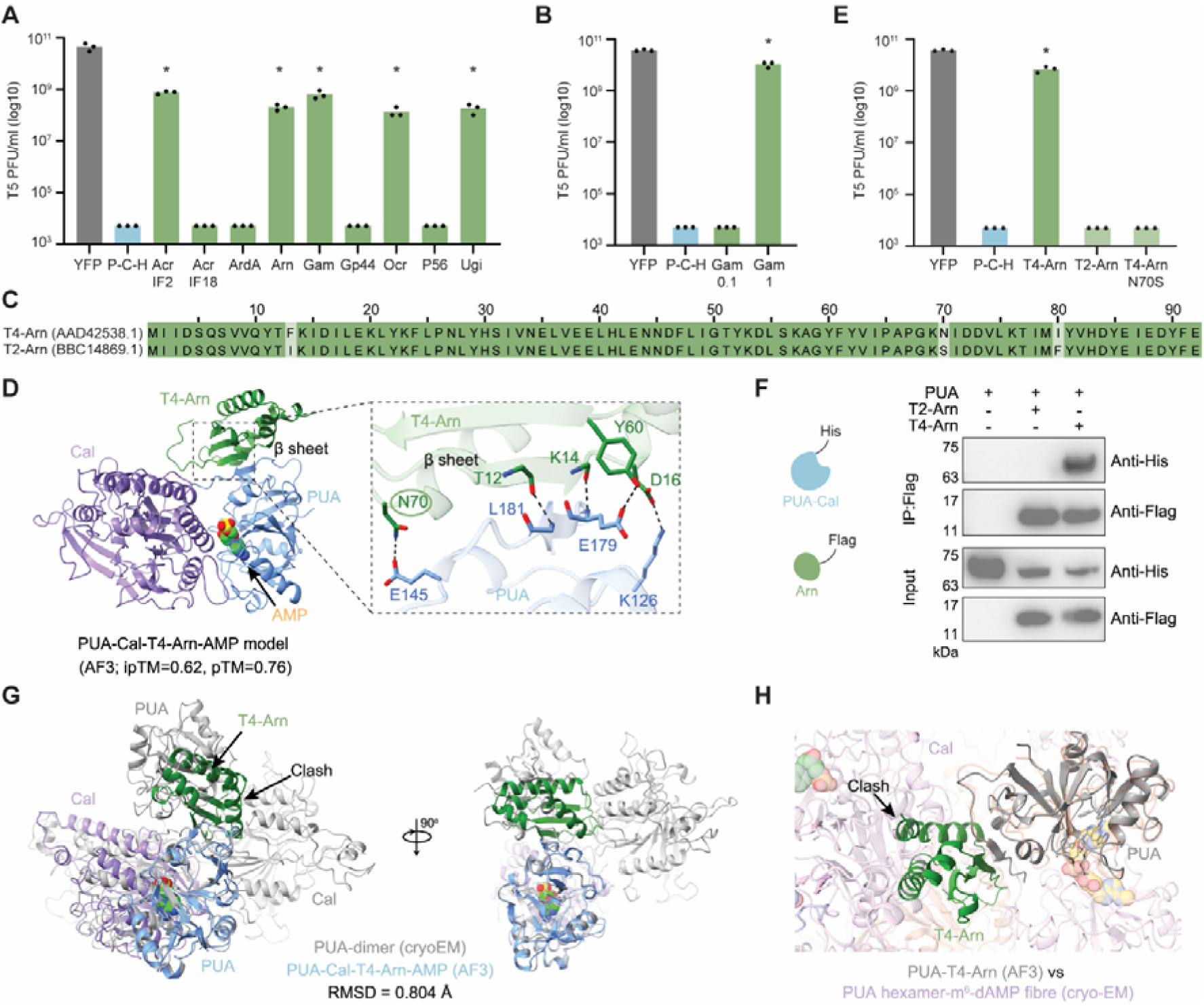
DNA mimics inhibit the anti-phage activity of PUA-Cal-HAD. **(A)** Effect of heterologous expression of DNA mimics on PUA-Cal-HAD (P-C-H) activity against phage T5. Bars represent the average of triplicates, with individual data points overlaid. YFP, yellow fluorescent protein (control); Asterisks indicate statistically significant differences to P-C-H (p < 0.05). **(B)** Effect of expressing Gam under Anderson promoter at variable expression levels (0.1, 1) on PUA-Cal-HAD activity against phage T5. Bars represent the average of triplicates, with individual data points overlaid. Asterisks indicate statistically significant differences to P-C-H (p < 0.05). **(C)** Alignment of T4-Arn and T2-Arn protein sequences, highlighting conserved regions. The Genbank accession numbers of the proteins are shown between brackets. **(D)** AlphaFold 3 (AF3) model of the PUA-Cal-T4-Arn. Left, model showing T4-Arn bound to the PUA-like domain. Right, close-up view of the T4-Arn-PUA interface, highlighting contacts between a β-sheet region of T4-Arn and residues on the PUA surface. **(E)** Effect of T4-Arn, T2-Arn, and T4-Arn with mutation N70S on PUA-Cal-HAD activity against phage T5. Bars represent the average of triplicates, with individual data points overlaid. Asterisks indicate statistically significant differences to P-C-H (p < 0.05). **(F)** Co-immunoprecipitation assays showing interaction between PUA-Cal and T4-Arn. His-tagged PUA-Cal efficiently co-precipitates with Flag-tagged T4-Arn, but not T2-Arn. **(G)** Structural superposition of the PUA-Cal-T4-Arn AF3 model with the cryo-EM PUA-Cal dimer. RMSD, root mean square deviation. **(H)** Superposition of the PUA-T4-Arn AF3 model onto the m^6^-dAMP-bound PUA-Cal filament obtained by cryo-EM.

Interestingly, phage T4, which encodes Arn, was not targeted by PUA-Cal-HAD, whereas the closely related phage T2 was efficiently restricted despite also encoding an Arn homolog (Figure 2B). Sequence comparison revealed divergency at three positions, including residues F13, N70, and I80 (Figure 7C).

To gain insight into the inhibitory mechanism, we generated an AlphaFold3 model of the PUA-Cal-T4 Arn complex. The best-scoring model predicts that T4-Arn binds directly to the PUA-like domain (Figure 7D). The model further predicts a key hydrogen bond between T4-Arn residue N70 and PUA residue E145, whereas T2 carries a serine at this position (Figure 7C). We hypothesised that this single substitution contributes to the differential sensitivity of T2 and T4 to PUA-Cal-HAD. Consistent with this, introducing the N70S substitution into T4-Arn abolished its ability to inhibit PUA-Cal-HAD (Figure 7E).

We next used co-immunoprecipitation assays to confirm the direct interaction between T4-Arn and PUA-Cal predicted by the AF3 model. These assays showed that T4-Arn binds efficiently to PUA-Cal, whereas T2-Arn does not interact, indicating that inhibitory activity is specific to the T4 protein (Figure 7F).

Structural superposition of the AF3 PUA-Cal-T4-Arn model with the cryo-EM PUA-Cal dimer shows that T4-Arn binding is incompatible with formation of a productive basal dimer geometry and prevents the dimer from adopting the precise relative orientation required for progression to the hexameric and filamentous states (Figure 7G). In this configuration, T4-Arn does not disrupt the overall folding of either PUA or Cal domains but instead occupies critical PUA surfaces required to stabilise productive PUA-PUA and PUA-Cal interactions within higher-order assemblies. Similarly, when the AF3 PUA-Cal-T4-Arn model is superimposed onto the m^6^-dAMP induced filament, several steric clashes with neighbouring PUA and Cal domains are observed (Figure 7H), demonstrating that T4-Arn binding and filament formation are mutually exclusive. These clashes arise because T4-Arn occupies the same spatial region required for close packing of adjacent hexamers, thereby physically blocking filament-compatible PUA conformations. As a result, the T4-Arn bound PUA-Cal complex is trapped in a non-productive monomeric configuration that cannot support hexamerisation.

Together, these results show that DNA mimics encoded by mobile genetic elements can antagonise PUA-Cal-HAD. For T4-Arn, this is accomplished by perturbing the PUA-dependent assembly transition into a hexamer and filament required for effector activity, and natural variation in Arn modulates this antagonism by altering direct engagement of the PUA-like domain.

## Discussion

This work defines PUA-Calcineurin-CE–HAD as a tripartite antiphage defence module that couples epigenetic danger sensing to a supramolecular effector state that rapidly collapses deoxyribonucleotide pools. The system detects infection through a host-encoded methylation mark (Dam-derived m^6^-dAMP) that becomes available specifically when phages degrade the bacterial chromosome. Ligand binding switches a preassembled PUA-Calcineurin-CE–HAD complex into a polymerising effector filament that amplifies and sustains nucleotide destruction.

DNA methylation is widely used in bacterial immunity, most canonically as a self/non-self marker in restriction-modification and related pathways^46^. Phages, in turn, deploy DNA-mimic anti-restriction proteins to neutralise methylation-linked barriers, illustrating how central methylation is to host–phage conflict^47^. PUA-Cal–HAD uses methylation in a fundamentally different way. Rather than recognising modified DNA as a substrate or reading methylation patterns to discriminate self from non-self, it senses a soluble breakdown product (m^6^-dAMP) generated when infection drives host genome degradation. This logic makes the trigger conditional on damage (DNA breakdown) rather than merely on the presence of modified DNA. In that sense, Dam methylation functions as a pre-existing barcode on the host chromosome that is converted into a danger signal only during phage attack. This theme resonates with the emerging view that several conflict systems can be wired to sense modified nucleic acids/nucleotides, but our work provides a direct mechanistic example in which the modified nucleotide itself is the activating ligand^8^. A useful comparison is retron Ec86, in which a phage-encoded methyltransferase activates a filamentous effector assembly by methylating retron-derived DNA (msDNA), culminating in NAD hydrolysis^48^. In contrast, PUA-Cal–HAD responds to methylated host-derived dAMP released during chromosome degradation and executes dNTP depletion. Together, these systems suggest that methylation can be exploited as a trigger at different layers, such as DNA, nucleotides, and metabolic small molecules, while retaining specificity for infection contexts.

An evolutionary interpretation of this behaviour is suggested by the broader genomic landscape of related PUA-like systems. Our contextual network analysis indicates that this PUA-like family recurrently mixes and matches with diverse effector outputs, including DNA-targeting modules (e.g. HNH nucleases and McrBC-like components) as well as predicted nucleotide-depleting enzymes, and it is frequently linked to N^6^-adenine methyltransferases and other enzymes associated with modified-nucleotide biochemistry. These associations are consistent with an arms-race logic in which phage strategies that evade or disable methylation-linked host defences select for host sensors that recognise modified nucleobases or nucleotides and then couple that recognition to either direct nucleic-acid attack or abortive infection through metabolic shutdown via degradation of key small molecules. In that framing, the mechanism we describe is particularly striking because it leverages a common phage strategy (host genome degradation for nucleotide scavenging) as the upstream cue that generates the soluble activating ligand.

Our biochemical and structural data support a model in which the two enzymatic components execute a sequential dephosphorylation cascade. HAD acts as a dATP γ-phosphatase, converting dATPs to dADPs by hydrolysing the β-γ phosphoanhydride bond (releasing inorganic phosphate). Calcineurin-CE then converts dADPs to dAMPs by hydrolysing the α-β phosphoanhydride bond (again releasing inorganic phosphate).

Although HAD-family enzymes are classically Mg^2+^-dependent phosphomonoesterases, nucleotide-directed activities are well represented across the superfamily. In particular, recent CARF-HAD immune effectors were shown to dephosphorylate ATP to and dATP to ADP in vitro, supporting that HAD domains can function as nucleotide-directed (d)NTP phosphohydrolases in antiviral pathways^49^. In PUA-Cal-HAD, our data place the HAD enzyme as an upstream effector that initiates high-flux depletion by removing the terminal phosphate of dNTP substrates, thereby feeding Calcineurin-CE with dNTP for processive conversion to dNMPs.

Importantly, the NAD^+^ cleavage activity we observed for Calcineurin-CE *in vitro* is HAD-independent and HAD does not suppress or enhance NAD^+^ turnover. This argues against a general allosteric role for HAD in Calcineurin-CE catalysis and instead supports a division of labour in which HAD is required primarily for reactions that depend on formation and consumption of dNDP intermediates.

Calcineurin-CE domains are predicted to support diverse nucleotide-targeting chemistries across bacterial immune systems, and such effectors may display broader substrate target selectivity *in vitro* than is realised physiologically^8^. In our hands, Calcineurin-CE can cleave NAD^+^ to NMN and AMP under *in vitro* conditions and in the absence of HAD coupling, yet cellular NAD^+^ levels remain largely unchanged upon defence activation, whereas dNTP pools collapse. We therefore interpret NAD cleavage as a latent Cal activity that is revealed when the effector is studied outside its physiological assembly/stoichiometry context, rather than as the primary effector output of the native PUA-Cal-HAD system. Several non-mutually-exclusive constraints could reconcile these observations. First, trigger availability and localisation likely bias the effector towards the dNTP pool: m^6^-dAMP binding to PUA is required to activate polymerisation, and that activated state may preferentially engage dNTP substrates (by geometry, access, or kinetics) while leaving NAD^+^ largely untouched *in vivo*. Second, stoichiometry and assembly may tune specificity: because HAD is preassembled with Calcineurin-CE in the native complex, physiological activation is wired for concerted dNTP to dNDP to dNMP conversion, whereas isolated Calcineurin-CE, or complexes formed at non-native ratios, may reveal alternative activities *in vitro*. Third, intracellular substrate competition and NAD(H) homeostasis could buffer NAD levels^50,51^, whereas dNTP pools are comparatively small and tightly regulated^52,53^, making them more vulnerable to catastrophic depletion once a dedicated sink is engaged.

An emerging theme in both bacterial and mammalian innate immunity is the assembly of higher-order supramolecular complexes that act as molecular switches and amplifiers. In mammals, viral RNA sensing triggers prion-like MAVS filaments to commit signalling to an all-or-none response, and cyclic dinucleotide binding promotes STING self-association into oligomeric assemblies that amplify downstream activation^54,55^. In bacteria, an expanding set of defence systems likewise relies on activation-coupled polymerisation. For example, bacterial TIR-STING and TIR-SAVED effectors form helical filaments/solenoids in response to cyclic nucleotide signals to activate TIR NADase activity and drive NAD⁺ depletion^56,57^, and the type III CRISPR-associated effector Cat1 forms filament networks that degrade NAD⁺ during antiviral defence^58^. More recently, invader nucleic acids have been shown to trigger filament formation in short Argonaute-associated systems (e.g., SPARDA/SPARHA), where polymerisation activates collateral nuclease effectors^59,60^, and filamentation can also activate bacterial NLR-like proteins such as Avs5^61^. PUA–Cal–HAD extends this paradigm in several distinctive ways. First, its trigger is a methylome-derived modified nucleotide (m^6^-dAMP) released by phage-driven degradation of the host chromosome, rather than a cyclic nucleotide second messenger or a nucleic-acid ligand. Second, the effector is preassembled: a Calcineurin-CE hexamer is already loaded with six HAD molecules before polymerisation, suggesting that filamentation primarily amplifies and stabilises preformed catalytic units rather than assembling an active site *de novo*. Third, its dominant physiological output is dNTP depletion, rather than NAD depletion or collateral nucleic-acid cleavage. Together, these features define a filament-forming immune strategy that links an epigenetic danger cue to rapid collapse of deoxyribonucleotide pools.

The emphasis on regulated assembly also highlights that stoichiometry and expression balance can be decisive for physiological mechanisms. In Gabija, for example, productive defence requires an appropriate GajA:GajB balance, underscoring that supramolecular immune machines can be stoichiometrically regulated rather than simply on/off^62^. More broadly, increasing defence system expression can broaden protection but frequently incurs a fitness cost or overt autoimmunity, highlighting how overexpression can confound mechanistic inference by pushing systems into non-physiological regimes^63^. In this context, both our observations (**Figure S7J**) and a recent inducible-expression study^64^ showing that PUA-Cal alone can protect are most parsimoniously interpreted as non-physiological regimes in which expression balance and stoichiometry are perturbed. Under such conditions, excess PUA-Cal may escape normal HAD coupling and regulated assembly, exposing secondary Calcineurin-CE activities. Consistent with this, the inducible-expression study reported NAD^+^-directed activity and NAD^+^ depletion^64^, whereas in the native locus we observe dominant dNTP depletion with minimal change in NAD^+^, suggesting that Calcineurin-CE substrate utilisation is tuneable by assembly state and component ratios. These comparisons emphasise the value of native-context analysis for defining physiological effector logic, while also pointing to potential applications of engineered or naturally HAD-lacking variants for controllable nucleotide/cofactor depletion.

Finally, the sensitivity of PUA-Cal–HAD to DNA mimic proteins highlights a familiar but underappreciated layer of conflict: phages and other mobile genetic elements frequently deploy DNA mimics such as Arn and Ocr to disable methylation-linked host defences^36,65,66^. The observation that closely related mimics (T4-Arn versus T2-Arn) differ in inhibitory potency suggests that anti-defence activity is evolvable but constrained, potentially by requirements imposed by other phage functions, by interactions with additional host pathways, or by trade-offs between broad antagonism and fitness. This fits a general picture in which phages carry modular anti-defence toolkits, but optimisation against one host system may be limited by pleiotropy and ecological context.

Together, these findings establish PUA-Cal–HAD as a defence system that (i) converts a host epigenetic mark into a ligand released specifically during phage-driven chromosome degradation, (ii) uses that ligand to switch a preassembled complex into a polymeric effector state, and (iii) executes immunity through rapid dNTP pool collapse driven by a coupled two-enzyme dephosphorylation circuit. More broadly, the contextual diversity of PUA-like defence loci suggests that modified-nucleotide sensing is repeatedly coupled to distinct effector logics across bacterial lineages, and that additional mechanisms linking modified-base recognition to nucleotide depletion likely remain to be discovered.

## Acknowledgements

We thank Alexander Harms (ETH Zurich) for providing the Basel phage collection, and Artem Isaev (Skolkovo Institute of Science and Technology) for providing the AcrIF2 and AcrIF18 plasmids. We also thank J. De La Cruz (MSK) and Sen Sagnik (MSK) for support with cryo-EM data collection. Y.W. is supported by a NIHR Southampton Biomedical Research Centre (BRC) Postdoctoral Bridging Fellowship. F.L.N. is supported by a Wessex Health Partners (WHP) and National Institute for Health and Care Research Wessex Experimental Medicine Network (NIHR WEMN) seed fund. D.J.P. is supported by the National Institutes of Health (NIH) grant GM145888, the Maloris Foundation, and the Memorial Sloan Kettering Cancer Center Core Grant (P30-CA008748). In addition to the MSKCC cryo-EM facilities, portions of this work were performed at the National Center for CryoEM Access and Training (NCCAT) and the Simons Electron Microscopy Center at the New York Structural Biology Center. These facilities are supported by the NIH Common Fund Transformative High Resolution Cryo-Electron Microscopy Program (U24 GM129539), the Simons Foundation (SF349247), and the New York State Assembly. G.G.N. and L.A. are supported by the Intramural Research Program of the NIH, National Library of Medicine (NLM). The contributions of the NIH authors are considered works of the United States Government. The findings and conclusions in this paper are those of the authors and do not necessarily reflect the views of the NIH or the U.S. Department of Health and Human Services.

## Author contributions

Conceptualization, F.L.N., D.J.P.; Methodology, F.L.N., D.J.P., Z.Z., Y.W., Y.J.L.; Investigation, Z.Z., Y.W., G.G.N., A.C., H.F., D.T., L.A.; Visualisation, Z.Z., Y.W., G.G.N., Y.J.L.; Writing – Original Draft, F.L.N., D.J.P., Z.Z., Y.W., G.G.N., L.A.; Writing – Review & Editing: all authors; Resources, F.L.N., D.J.P., P.W.; Funding Acquisition, F.L.N., D.J.P., L.A., P.W.

## Methods

### Materials availability

All unique bacterial strains, phages, and plasmids generated in this study are available from the lead contact without restriction. Raw data have been deposited at Zenodo (https://zenodo.org/records/18262648?preview=1&token=eyJhbGciOiJIUzUxMiJ9.eyJpZCI6ImJhNWRhNzJjLTQ0MjctNDU3MC04YzU4LWIwOTcwYzRlNTdhNyIsImRhdGEiOnt9LCJyYW5kb20iOiJkNzgyNzIxZWI0OTU3Njg0ZGVmNjA3MGQzMDU5NjVlMiJ9.Kw2IFrmY1QQld8DifVARKxcB1aJpSVQXGTBdttpY93wNMw1aBJKvh_t6zkg5Qtvwh0LZUzIt_MteJJ5iJlU13w), Protein Data Base (PDB: 9ZWP, 9ZXB, 9ZXE, 9ZXF, 9ZXG, 9ZXH, 9ZXI, 10CV) and the Electron Microscopy Data Bank (EMDB: EMD-74917, EMD-74918, EMD-74921, EMD-74922, EMD-74923, EMD-74924, EMD-74925, EMD-75078) and are publicly available as of the date of publication.

### Bacterial strains and phages

*E. coli* strains (**Table S4**) were grown at 37 °C in Lysogeny Broth (LB) for liquid cultures or LB agar (LBA, 1.5 % (w/v) agar) for solid cultures. Whenever applicable, LB was supplemented with 25 µg/ml of chloramphenicol, 100 µg/ml of ampicillin, or 50 µg/ml of kanamycin. Phages used in this study are described in **Table S4**, and phage infection was performed at 37 °C.

### Sequence analysis

Sequence similarity searches were conducted using PSI-BLAST^67^ and JackHMMER^68^ against the NCBI non-redundant protein database (nr) or a version clustered down to 50% sequence identity (nr50). The searches were initiated using the previously identified representatives of each family of domains considered in this work. Sequence-similarity-based clustering (percentage identity or bit-score) was performed using the MMseqs program^69^, with the clustering parameters adjusted according to specific goals, such as redundancy removal, definition of homologous groups, and creation of new profiles. Multiple sequence alignments (MSA) were generated using the MAFFT program^70^ with the local-pair algorithm, combined with the parameters –maxiterate 3000, –op 1.5, and –ep 0.2, and were manually refined based on structural superpositions and profile-profile comparisons.

### Comparative genomics, domain identification, and phylogenetic analysis

Genomic neighbourhoods were extracted from genomes available in the NCBI Genome database using scripts written in Perl and Python (https://zenodo.org/records/13864123). Further analysis of these genomic neighbourhoods was performed by clustering the protein products of neighbouring genes. Domains were then identified using a collection of HMMs and PSSMs maintained by the Aravind lab, along with HMMs from the Pfam database^71^, via the RPSBLAST and HMMSCAN programs. To further refine detection, domain identification was extended through remote homology analysis using HHpred^72^ against profiles built from the Pfam and PDB70^73^ databases.

### Contextual network construction

The domain architectures and conserved gene neighbourhoods were decomposed into their constituent domains. The contextual connections were then converted into an order graph with the domains as its nodes and the adjacency relationships as edges. The communities of domains were identified within the graph using the Louvain algorithm^74^, and the graph was rendered using the Fruchterman-Reingold^75^ followed by the Kamada-Kawai algorithm^76^. Network analysis was performed using the functions of the R igraph^76,77^ or Python networkX libraries^78^.

### Cloning

Plasmids and primers used in this work can be found in **Tables S5** and **S6**, respectively. PUA-Cal-HAD from *E. coli* ECOR28 was cloned into pACYC-duet by Gibson assembly. For protein expression and purification, PUA-Cal was cloned into the pET21a vector to generate a C-terminal His-tagged fusion protein and HAD was cloned into pYB100 vectors carrying either a C-terminal His tag or a C-terminal flag tag. DNA mimic genes were cloned into pCOLA-duet by Gibson assembly. The plasmids were recovered in DH10B cells, extracted using Monarch Plasmid Miniprep Kit (New England Biolabs) and confirmed by sequencing at Macrogen (Netherlands) or Plasmidsaurus (USA). Mutations of PUA-Cal-HAD were engineered by around-the-horn PCR or Gibson assembly and confirmed by Sanger sequencing at Macrogen. Plasmids were transformed individually or in combinations into competent BL21-AI cells prepared using the Mix&Go! *E. coli* Transformation Kit (Zymo Research).

### Phage infectivity

Overnight bacterial cultures were mixed with 0.6% top agar and overlaid on LBA plates. For cultures containing inducible DNA mimic-expressing plasmids, overnight cultures of the bacteria were diluted 1:50 in LB containing antibiotics, induced with 0.2% L-arabinose, and incubated for 4 hours before being used in double agar overlay assays. Ten-fold serial dilutions of the phage stocks were spotted onto the bacterial lawn and the plates incubated overnight at 37 °C. The phage plaques were counted and used to calculate the efficiency of plating relative to the control. Statistical significance was determined using the multiple comparison function from Two-way ANOVA with a p-value of <0.01.

### Liquid assay

Overnight bacterial cultures were diluted to an optical density at 600 nm (OD_600_) of 0.1 in LB containing antibiotics and 0.2% arabinose when needed. The bacterial suspension was distributed into wells of a 96-well plate to which phage dilutions or LB were added. The plates were incubated in a Clariostar Plus plate reader at 37 °C with shaking at 200 rpm, with OD_600_ measured every 5 min for 24 h.

### Time post infection assay

Overnight bacterial cultures were diluted to an optical density at 600 nm of 0.1 in LB containing antibiotics and 0.2% arabinose when needed. The cultures were infected with phage at an MOI of 10 or 0.01. At 0-, 15-, 30-, 45- and 60-minutes post infection, a sample was taken and centrifuged at 12,000 × *g* for 2 minutes. The supernatant was serially diluted, and the phages were quantified by plaque assay on a bacterial lawn of cells with YFP. PFUs were counted after overnight incubation at 37 °C. To determine the CFUs, the pellets were washed twice with LB, the cells resuspended and serially diluted. The dilutions were spotted onto LBA plates supplemented with antibiotics, and the plates incubated overnight at 37 °C.

### Generation of phage escape mutants

Phage plaques obtained in double layer agar plates with PUA-Cal-HAD were picked and dissolved in 30 µl of LB. The recovered phages were produced in liquid culture using the PUA-Cal-HAD-expressing strain as the host. The produced phages were serially diluted and spotted onto bacterial laws of PUA-Cal-HAD-expressing and control cells, to identify those phages able to escape PUA-Cal-HAD native defence.

### Phage DNA extraction and sequencing

Phage DNA was extracted using the phenol-chloroform method^6^. Briefly, DNase I and RNase were added to the phage stock at 1 µg/ml each, and the solution was incubated for 30 min. EDTA at 20 mM, Proteinase K at 50 µg/ml, and SDS at 0.5% were added to the solution and incubated for 1 hour at 56 °C. An equal volume of phenol was added to the mixture, vortexed, and centrifuged at 3,000 × *g* for 10 min, at room temperature. The aqueous phase was recovered, mixed with an equal volume of a 1:1 mixture of phenol:chloroform, and centrifuged. The procedure was repeated with chloroform. After centrifugation, the aqueous phase was recovered and mixed with 0.1 volumes of 3 M sodium acetate, pH 5.0 and 2.5 volumes of ice-cold absolute ethanol. The mixture was incubated at −20 °C for 1 hour and centrifuged at 14,000 × *g* for 15 min. The DNA pellet was resuspended in ice-cold 70% ethanol, centrifuged for 5 min, and let dry at room temperature before resuspending in nuclease-free water. Phage samples were sequenced at Novogene (UK).

### Analysis of phage escape mutants

For wild-type phage, seqtk version 1.3 (https://github.com/lh3/seqtk) was used to sample reads at 100x depth of the phage genomes. The subset reads were assembled using SPAdes version v3.15.3^79^ using default settings. For phage escape mutants, all reads were mapped to the assembled wild-type phage genome using bwa version 0.7.17^80^ and samtools version 1.11^81^. SNP calling was performed using gatk version 4.2.6.1^82^.

### LC-MS profiling of bacterial metabolites

Overnight bacterial cultures were diluted to an OD_600_ of 0.1 in LB containing antibiotics. The cells were incubated at 37 °C, 200 rpm until an OD_600_ of 0.6. Phage T2 was added at an MOI of 10, while control cells remained uninfected. After 15 min of incubation at 37 °C, 200 rpm, cells were spun down at 4,000 × *g* for 10 min and the pellets frozen in liquid nitrogen. Cell metabolites were extracted by dissolving the pellets in 2 mL of 100 mM sodium phosphate buffer, pH 7.5. Samples were lysed by sonication at 4°C using a 2 s on and 5 s off cycle at 10% amplitude. The lysed samples were centrifuged at 15,000 × *g* for 30 min at 4°C, and the supernatant was concentrated on an Amicon Ultra-0.5 Centrifugal Filter Unit 3LJkDa (Merck Millipore) by centrifuging 15,000 × *g* for 1 hour at 4°C. The lysates were then analysed by LC-MS. Polar metabolites were profiled using a Waters Acquity Premier HSS T3, 1.8um, 2.1×100mm column in an Acquity Premier UPLC-MS System (Thermo Fisher Scientific), operated in positive ionization mode. LC was carried out using a gradient of 1% to 10% acetonitrile (mobile phase A: water with 0.05% TFA; mobile phase B: acetonitrile). The total run time was 8 min with a gradient between 1 to 5 min. m^6^-dAMP was prepared by digesting 10 mM m^6^-dATP with Apyrase (New England Biolabs) for 1h at 37°C, following manufacturer instructions, followed by filtration with an Amicon Ultra-0.5 Centrifugal Filter Unit 3 kDa. The flow-through was injected in Waters LC Prep system with a Waters XBridge BEH C18 OBD Prep Column (5 µM, 19×150 mm, buffers and gradient as above), with collection of the m/z 346.10 peak.

### Protein expression

For protein expression, *E. coli* BL21 (DE3) cells were transformed with the expression plasmids and grown in LB supplemented with antibiotics at 37 °C to an OD_600_ of 0.6-0.8. For PUA-Cal-HAD co-expression experiments, ampicillin-resistant PUA-Cal-His and kanamycin-resistant HAD-Flag plasmids were co-transformed into *E. coli* BL21 (DE3) cells and cultured in the presence of both antibiotics. Protein expression was induced with 0.5 mM Isopropyl β-D-1-thiogalactopyranoside (IPTG), and cultures were incubated at 18 °C for 16-18 h. Cells were harvested by centrifugation at 4,000 × *g* for 15 min at 4 °C and stored at −80 °C.

### Purification of PUA-Cal

Cell pellets were resuspended in lysis buffer containing 25 mM Tris-HCl (pH 8.0), 300 mM NaCl, 2% (v/v) glycerol, and 1 mM DTT, supplemented with cOmplete EDTA-free protease inhibitor cocktail, and lysed by sonication on ice. Lysates were clarified by centrifugation at 22,000 rpm for 1 hour at 4 °C. The supernatant was applied to Ni-NTA affinity resin equilibrated in lysis buffer. After extensive washing with buffer containing 25 mM Tris-HCl (pH 8.0), 300 mM NaCl, and 30 mM imidazole, bound proteins were eluted with buffer containing 500 mM imidazole. Eluted PUA-His was further purified by SEC using a Superdex 200 Increase 10/300 GL column (GE Healthcare) equilibrated in SEC buffer containing 25 mM HEPES (pH 7.5), 150 mM NaCl, 0.2% (v/v) glycerol, and 1 mM DTT. PUA-His eluted as three distinct species corresponding to monomeric, dimeric, and hexameric assemblies. Each oligomeric fraction was collected separately, concentrated individually to approximately 8 mg/mL, flash-frozen in liquid nitrogen, and stored at −80 °C. For purification of PUA-Cal-His under metal-ion conditions, the protocol described above was used with the exception that all buffers (from lysis through SEC) were supplemented with 10 mM MgCl₂, 5 mM FeCl_3_, and 2 mM MnCl₂.

### Purification of HAD

HAD-His was expressed and purified using the protocol described for PUA-Cal-His. Final SEC was performed using a Superdex 75 Increase 10/300 GL column (GE Healthcare). HAD-His eluted as a monomeric species, and purity was confirmed by SDS-PAGE. Due to protein precipitation and degradation at higher concentrations, HAD-His was concentrated to 1.8 mg/mL, flash-frozen in liquid nitrogen, and stored at −80 °C prior to use. For HAD-Flag purification, cell pellets were resuspended in the same lysis buffer described above and lysed by sonication. Lysates were clarified by centrifugation at 40,000 rpm for 1 h at 4 °C. The supernatant was incubated with anti-FLAG M2 affinity resin (Sigma) for 3 h at 4 °C with gentle mixing. The resin was washed extensively with lysis buffer, and bound proteins were eluted using lysis buffer supplemented with 0.2 mg/mL 3×FLAG peptide. Eluted proteins were concentrated and further purified by SEC using a Superdex 200 Increase 10/300 GL column (GE Healthcare) equilibrated in SEC buffer (25 mM HEPES (pH 7.5), 150 mM NaCl, 0.2% glycerol, and 1 mM DTT). HAD-Flag eluted as a monomeric species, and purity was confirmed by SDS-PAGE.

### Expression and purification of the PUA-Cal-HAD complex

For co-expression of the PUA-Cal-HAD complex, *E. coli* BL21 (DE3) cells co-transformed with PUA-Cal-His and HAD-Flag plasmids were induced and harvested under the same conditions described above. Co-expressed complexes were purified using anti-Flag affinity resin following the same protocol used for HAD-Flag purification and the fractions obtained were analysed by SDS-PAGE. The eluted complex was concentrated and subjected to SEC using a Superdex 200 Increase 10/300 GL column (GE Healthcare) to assess oligomerisation. Most of the protein eluted as a high-molecular-weight aggregate (peak 1), as confirmed by SDS-PAGE analysis. Fractions corresponding to peak 2 were immediately reloaded onto the same column, however, the protein continued to aggregate during the second run. These results indicate that the PUA-Cal-HAD complex is unstable in solution and prone to aggregation under the conditions tested.

### Cryo-EM sample preparation and data collection

Purified proteins or complexes were concentrated to 8 mg/mL. For PUA-Cal-His samples containing m^6^-AMP, m^6^-AMP was added to a final concentration of 0.5 mM in metal ion-supplemented buffer, and the sample was incubated on ice for 30 min prior to grid freezing. For PUA-Cal-His samples forming m^6^-dAMP-induced filaments, m^6^-dAMP was added to a final concentration of 0.5 mM in metal ion-supplemented buffer, and the sample was incubated on ice for 5 min only and vitrified immediately before the onset of protein precipitation. PUA-Cal-His dimeric and hexameric samples were vitrified at a protein concentration of 8 mg/mL without prior incubation. To mitigate preferred orientation, fluorinated octyl maltoside (FOM) was freshly added to all samples immediately prior to freezing to a final concentration of 0.5 mM. A 4 μL aliquot of each sample was applied to glow-discharged holey carbon gold grids (Quantifoil Au 300 mesh R1.2/1.3). Grids were blotted for 2.5 s at 100% humidity and 6 °C, then plunge-frozen into liquid ethane using a Vitrobot Mark IV (FEI). Grid screening was performed on a 300 kV Titan Krios G3 transmission electron microscope equipped with a K3 direct electron detector and operated using SerialEM software (Thermo Fisher Scientific).

For the PUA-Cal-His dimer sample and the PUA-Cal-His hexamer sample in the presence of m^6^-AMP or m^6^-dAMP, cryo-EM data were collected at the Memorial Sloan Kettering Cancer Center (MSKCC) using a Titan Krios G2 transmission electron microscope (FEI) operated at 300 kV and equipped with a K3 direct electron detector. Data acquisition was controlled using SerialEM software. Movies were recorded in super-resolution mode with a total electron dose of 53 e⁻ Å^-2^, a defocus range of −0.8 to −2.2 µm, and a calibrated pixel size of 1.064 Å.

For the PUA-Cal-His hexamer and the PUA-Cal-His hexamer in the presence of metal ions, data were collected at MSKCC using a Titan Krios G4 transmission electron microscope (FEI) operated at 300 kV and equipped with a Falcon 4i electron detector. Data collection was performed using EPU software. Movies were recorded in counting mode with 45 EER frames. The defocus range was set from −0.8 to −2.2 µm, with a pixel size of 0.725 Å and a total electron dose of 60 e⁻ Å^-2^.

### Cryo-EM data processing

All cryo-EM data collection and refinement statistics are detailed in **Table S7**. For PUA-Cal dimer structure determination, a total of 6,563 movies were processed using cryoSPARC (v4.2.1)^83^. Patch-based motion correction and patch-based contrast transfer function (CTF) estimation were applied to correct beam-induced motion and estimate CTF parameters, respectively. Micrographs exhibiting ice contamination, high astigmatism, or poor CTF-fit resolution were excluded by applying appropriate threshold criteria using the *Manually Curate Exposures* job. The remaining high-quality micrographs were subjected to particle picking using the Blob Picker, followed by two-dimensional (2D) classification. Representative high-quality classes were selected to generate templates for subsequent template-based particle picking. Particles were then extracted, yielding a total of 3,302,016 particles. Multiple rounds of 2D classification were performed to remove contaminants and poorly aligned particles. Following *ab initio* reconstruction and heterogeneous refinement into three classes, one well-resolved class was selected for subsequent homogeneous refinement and non-uniform refinement, yielding a reconstruction at 3.43 Å resolution based on 820,273 particles. These particles were subsequently subjected to global and local CTF refinement to correct residual astigmatism and higher-order aberrations, followed by a final round of non-uniform refinement. This improved the overall resolution to 3.24 Å and substantially enhanced map quality.

For PUA-Cal hexamer structure determination, a total of 9,199 movies was processed using cryoSPARC. Patch-based motion correction and patch-based CTF estimation were applied to correct beam-induced motion and estimate CTF parameters. Micrographs were then subjected to particle picking using the Blob Picker followed by particle extraction, yielding a total of 1,918,576 particles. Multiple rounds of 2D classification were performed to remove contaminants and poorly aligned particles. Following two rounds of *ab initio* reconstruction and heterogeneous refinement into three classes, two distinct hexameric conformational states were identified: state 1 (two ‘up’ and one ‘down’ dimers) and state 2 (three ‘down’ dimers), both in the presence of AMP. These states contained 110,488 and 100,573 particles, respectively. Each state was further refined by homogeneous refinement, followed by global and local CTF refinement to correct higher-order aberrations. Final non-uniform refinement yielded reconstructions at 2.91 Å (state 1) and 2.95 Å (state 2) resolution.

For PUA-Cal hexamer with metal ions bound structure determination, a total of 9,001 movies were processed using cryoSPARC. Beam-induced motion was corrected using patch-based motion correction, and CTF parameters were estimated using patch-based CTF estimation. Particle picking was initially performed using the Blob Picker, followed by 2D classification. Representative high-quality classes were selected to generate templates for subsequent template-based particle picking, after which particles were extracted, yielding a total of 1,537,901 particles. Two additional rounds of 2D classification were carried out to remove contaminants and poorly aligned particles. Three-dimensional classification resolved two distinct hexameric conformational states in the presence of AMP: state 1, in which each dimer adopts a configuration with two protomers in the ‘up’ conformation and one in the ‘down’ conformation, and state 2, in which all three protomers adopt the ‘down’ conformation. Particles corresponding to these two well-resolved states were selected for further processing and refined independently by homogeneous refinement. Global and local CTF refinement were then applied to improve optical parameter estimation, followed by a final round of non-uniform refinement. This yielded reconstructions at 2.67 Å resolution based on 512,205 particles for state 1 and 2.76 Å resolution based on 204,473 particles for state 2.

For the m^6^-AMP-bound PUA-Cal hexamer structure determination, a total of 4,433 movies were processed in cryoSPARC, beam-induced motion correction and CTF estimation were performed using patch-based workflows. Particle picking was carried out with the Blob Picker, followed by particle extraction, yielding 2,615,296 particles. Two iterative rounds of 2D classification were applied to remove contaminants and poorly aligned particles. *Ab initio* reconstruction and heterogeneous refinement, performed in two successive rounds, resolved two predominant hexameric conformations in the presence of m^6^-AMP, as observed for AMP: state 1 and state 2. These states comprised 398,253 and 199,430 particles, respectively. Each class was refined independently by homogeneous refinement, followed by global and local CTF refinement, and a final round of non-uniform refinement yielded reconstructions at 2.95 Å (state 1) and 3.21 Å (state 2) resolution.

For the m^6^-dAMP-bound PUA-Cal fibre structure determination, a total of 6,903 movies were processed using cryoSPARC. Patch-based motion correction and patch-based CTF estimation were performed to correct beam-induced motion and estimate CTF parameters. Micrographs exhibiting ice contamination, high astigmatism, or poor CTF-fit resolution were removed using threshold-based filtering with the *Manually Curate Exposures* job. The remaining high-quality micrographs were subjected to particle picking using the Manual Picker to select filamentous particles, followed by 2D classification. Representative high-quality 2D classes were chosen to generate templates for subsequent template-based particle picking. Particles were then extracted with binning by a factor of two (bin2), yielding a total of 208,369 particles. Several additional rounds of 2D classification were performed to further remove contaminants and poorly aligned particles. Following *ab initio* reconstruction and heterogeneous refinement into four classes, a single well-resolved class containing 32,491 particles was selected for further processing. These particles were refined by homogeneous refinement and helical refinement. For higher-resolution refinement, particles were re-extracted at full pixel size (bin1), and the corresponding bin2 volume was rescaled and transferred to bin1 using the RELION handler in RELION^84^. The particles were subsequently subjected to global and local CTF refinement to correct residual astigmatism and higher-order aberrations, followed by a final round of non-uniform refinement using the bin1 volume. This processing strategy improved the overall reconstruction to a resolution of 3.09 Å based on 32,186 particles, substantially enhanced map quality.

### Model building and refinement

Initial atomic models were generated using AlphaFold 3^85^ predictions and docked into cryo-EM density maps. Manual model building was performed in Coot^86^ to adjust side chains, loops, and ligand positions. Real-space refinement was carried out in PHENIX^87^ with geometry restraints. Model quality was assessed using standard validation metrics, including Ramachandran statistics and map-to-model correlation.

### Structural alignment, analysis and visualisation

Structural alignments and root-mean-square deviation (RMSD) calculations were performed using UCSF ChimeraX^88^. Alignments were carried out using either the PUA-like domains or the Cal domains, as indicated. Structural figures were prepared using UCSF ChimeraX and UCSF Chimera^89^, and composite figures were assembled using Adobe Photoshop and Adobe Illustrator.

### Co-immunoprecipitation assays for PUA-Cal-HAD and PUA-Cal-Arn complex formation

Flag-tagged HAD, T2-Arn or T4-Arn expression plasmids were co-expressed with the His-tagged PUA-Cal expression plasmid. Cells were cultured in the presence of antibiotics, and protein induction and cell harvest were performed using the same protocol described above. PUA-Cal-His expressed alone was used as a negative control. Protein purification was carried out as described above using anti-FLAG affinity resin, except that 0.2% (v/v) Triton X-100 was included from the lysis step onward throughout the purification to reduce non-specific binding. The resin was washed thoroughly five times with wash buffer containing Triton X-100 prior to elution with 3×FLAG peptide. Elution was performed on ice for 30 min with gentle mixing every 5 min. Eluted proteins were subjected to SDS-PAGE. Proteins were electro-transferred onto PVDF membranes using semi-dry transfer cell (Bio-Rad). PVDF membranes were blocked using 5% (w/v) non-fat dry milk (Bio-Rad) for 1h at room temperature. Following incubation with horseradish peroxidase (HRP)-conjugated anti-Flag tag (86861S, Cell Signaling) or anti-His tag antibodies (MA1-21315-HRP, Thermo Fisher) (1:2,000-fold dilution (v/v)) overnight at 4°C, PVDF membranes were washed in 0.05% Tween-20/TBS. The bound antibodies were visualised by using an enhanced chemiluminescence reagent (Thermo Scientific) and quantified by densitometry using a ChemiDoc MP imaging system (Bio-Rad).

### Pull down assay for PUA-Cal-HAD complex formation

To assess direct interactions between PUA-Cal and HAD, affinity pull-down assays were performed using Flag-tagged HAD as described above. Briefly, 15 μg of HAD-Flag monomer was incubated separately with 15 μg of either PUA-Cal-His dimer or PUA-Cal-His hexamer and diluted to a final volume of 500 μL in SEC buffer supplemented with 0.2% (v/v) Triton X-100 to reduce non-specific interactions. As negative controls, 15 μg of PUA-Cal-His dimer or PUA-Cal-His hexamer alone were processed in parallel. Samples were rotated for 30 min at 4 °C prior to the addition of anti-FLAG affinity resin. After incubation, 20 μL of each sample were reserved as input, and the remaining volume was incubated with pre-equilibrated anti-Flag resin for 2 h at 4 °C with gentle rotation. The resin was washed five times with SEC buffer containing 0.2% Triton X-100 and eluted with the same buffer supplemented with 0.2 mg/mL 3×Flag peptide. Eluted proteins were analysed by western blotting using HRP-conjugated antibodies against the Flag tag and the His tag at a 1:2,000 (v/v) dilution to detect HAD and PUA, respectively. Bound antibodies were visualised using enhanced chemiluminescence reagents, and signal intensities were quantified by densitometry using a ChemiDoc MP imaging system (Bio-Rad).

### Electrophoretic mobility shift assays

For binding assays assessing interactions between PUA dimers or hexamers and DNA substrates, single-stranded and double-stranded DNA containing m^5^C or m^6^A modifications, as well as unmodified controls, were used. The oligonucleotides and modified oligonucleotides were synthesised by Integrated DNA Technology (IDT). The m^6^A-containing ssDNA substrate consisted of a 15-nt sequence (5′-ATATCGT[m6-dA]GGTATGG-3′) together with its complementary strand. The m^5^C-containing ssDNA substrate consisted of a 15-nt sequence (5′-ATATCGT[m5-dC]GGTATGG-3′) and its complementary strand. An unmodified 15-nt DNA sequence (5′-ATATCGTAGGTATGG-3′) and its complementary strand were used as controls. Double-stranded DNA substrates were generated by self-annealing complementary oligonucleotides in annealing buffer containing 25 mM HEPES (pH 7.5), 150 mM NaCl, and 2 mM MgCl₂. DNA substrates were used at a final concentration of 10 nM in a total reaction volume of 20 μL. The molar ratios of PUA-Cal hexamer or PUA-Cal dimer to DNA substrate were set at 0, 1, 8, 16, 32, 64, and 128:1. Protein and DNA were diluted in annealing buffer supplemented with an additional 7.5% (v/v) glycerol to facilitate gel loading. Binding reactions were incubated on ice for 1 h, after which 10 μL of each sample were loaded onto 8–20% 15-well TBE polyacrylamide gels (Invitrogen). Electrophoresis was performed at 100 V for 80 min at 4 °C in 1× TBE buffer. Nucleic acids were visualized by staining with SYBR Gold nucleic acid gel stain (Invitrogen) for 30 min in the dark and imaged using a ChemiDoc imaging system.

### Mass photometry

Mass photometry experiments were performed using a Refeyn TwoMP instrument. A pre-assembled 6-well sample cassette (Refeyn) was mounted at the centre of a clean sample carrier slide (Refeyn), with each well used for an individual measurement. To establish the focal plane, 15 μl of freshly prepared SEC buffer (25 mM HEPES, pH 7.5, 150 mM NaCl, 0.2% (v/v) glycerol) or metal ion-supplemented buffer containing an additional 10 mM MgCl₂, 5 mM FeCl_3_, and 2 mM MnCl₂ in SEC buffer was added to each well as a blank. Focus was determined and maintained throughout data acquisition using the built-in autofocus system based on total internal reflection. Purified PUA-Cal dimers, PUA-Cal hexamers, HAD monomers, PUA-Cal dimer-HAD complexes, and PUA-Cal hexamer-HAD complexes, either alone or in the presence of m^6^A-containing ssDNA or m^6^-dAMP, were prepared under the same buffer conditions used during purification and incubated on ice for 5 min prior to measurement. Samples were initially diluted to 200 nM, and 3 μL of the diluted protein solution was added to the buffer drop in the measurement well, resulting in a final protein concentration of 33.3 nM. After autofocus stabilisation, movies were recorded for 60 s. Data acquisition was performed using Refeyn AcquireMP (v2024.1.1.0), and data analysis was carried out using Refeyn DiscoverMP (v2024.1.0.0). Contrast-to-mass calibration was performed using bovine serum albumin (BSA; Sigma) as a standard. Statistical analysis was conducted in DiscoverMP, where Gaussian fitting was applied to mass distribution peaks to determine the average molecular mass of each species. Plots were generated using GraphPad Prism 10.

### PUA-Cal-HAD *in vitro* activity on (deoxy)nucleotides and NAD^+^

NAD^+^, dNTPs, dNMPs, NTPs, or NMPs were prepared at a concentration of 0.1 mM in a buffer composed of 50 mM Tris-HCl (pH 7.5) and 1 mM MgCl_2_. m^6^-dAMP and m^6^-AMP were used as variables at a concentration of 10 µM. PUA-Cal, HAD or a combination of both, were added to each nucleotide solution mixture at a concentration of 1 µM. The reactions were carried out at 37°C for 1-3h. Thermolabile proteinase K (0.12 units) was added to each reaction and incubated at 25 °C for 4h before heating at 55°C for 15 min. The product was diluted with an equal volume of LC aqueous mobile phase (10 mM ammonium acetate, pH 4.5) and filtered through a 0.2 µm PTFE filter (Millipore). Half of the filtrate was used for LC-MS analysis as described below.

### LC-MS analysis of *in vitro* PUA-Cal-HAD enzymatic reactions

LC-MS analysis was conducted on an Agilent 1290 Infinity II UHPLC-MS system equipped with a G7117 Diode Array Detector and connected to an Agilent 6135 LC/MSD XT Single Quadrupole Mass Detector. Reverse-phase liquid chromatography employed a Waters XSelect HSS T3 C18 column (4.6 × 150 mm, 3 µm particle size) operated at a flow rate of 0.6 ml/min with a binary linear gradient mobile phase consisting of 10 mM ammonium acetate (pH 4.5) and methanol. The course of chromatography was monitored at 260 nm. Mass spectrometry was operated in both positive (+ESI) and negative (-ESI) electrospray ionization modes. MS was performed with a capillary voltage of 2500 V in both modes, a fragmentor voltage of 70 V, and a mass range of m/z 100-1000. Agilent ChemStation software was used for the preliminary LC-MS data acquisition, analysis, and generation of LC-UV traces, MS spectra, and integration report. ChemStation-produced chromatograms were further annotated in Adobe Illustrator.

## QUANTIFICATION AND STATISTICAL ANALYSIS

Unless stated otherwise, experiments were performed in biological triplicates and are represented as the mean and standard deviation. Statistical significance was calculated by ratio paired t-test or by two-way ANOVA with sidak’s multiple comparison test, with a significance level of <0.05.

## References

1. Georjon, H., and Bernheim, A. (2023). The highly diverse antiphage defence systems of bacteria. Nat Rev Microbiol. 10.1038/s41579-023-00934-x.

2. Laub, M.T., and Typas, A. (2024). Principles of bacterial innate immunity against viruses. Curr Opin Immunol 89, 102445. 10.1016/j.coi.2024.102445.

3. Loeff, L., Walter, A., Rosalen, G.T., and Jinek, M. (2025). DNA end sensing and cleavage by the Shedu anti-phage defense system. Cell 188, 721–733.e717. 10.1016/j.cell.2024.11.030.

4. Yuping, L., Guan, L., Becher, I., Makarova, K.S., Cao, X., Hareendranath, S., Guan, J., Stein, F., Yang, S., Boergel, A., et al. (2025). Jumbo phage killer immune system targets early infection of nucleus-forming phages. Cell 188, 2127–2140.e2121. 10.1016/j.cell.2025.02.016.

5. Muralidharan, A., Costa, A.R., Fierlier, D., van den Berg, D.F., van den Bossche, H., Zoumaro-Djayoon, A.D., Pabst, M., Pacesa, M., Correia, B.E., and Brouns, S.J.J. (2025). Molecular basis for anti-jumbo phage immunity by AVAST Type 5. bioRxiv, 2025.2007.2008.663546. 10.1101/2025.07.08.663546.

6. Zhang, Z., Todeschini, T.C., Wu, Y., Kogay, R., Naji, A., Rodriguez, J.C., Mondi, R., Kaganovich, D., Taylor, D.W., Bravo, J.P.K., et al. (2025). Kiwa is a membrane-embedded defence supercomplex activated at phage attachment sites. Cell 188, p5862–5877.e23. 10.1016/j.cell.2025.07.002.

7. Kibby, E.M., Conte, A.N., Burroughs, A.M., Nagy, T.A., Vargas, J.A., Whalen, L.A., Aravind, L., and Whiteley, A.T. (2023). Bacterial NLR-related proteins protect against phage. Cell 186, 2410–2424.e2418. 10.1016/j.cell.2023.04.015.

8. Nicastro, G.G., Burroughs, A.M., Iyer Lakshminarayan M., and Aravind, L. (2023). Functionally comparable but evolutionarily distinct nucleotide-targeting effectors help identify conserved paradigms across diverse immune systems. Nucleic Acids Res 51, 11479–11503. 10.1093/nar/gkad879.

9. Payne, L.J., Hughes, T.C.D., Fineran, P.C., and Jackson, S.A. (2024). New antiviral defences are genetically embedded within prokaryotic immune systems. bioRxiv, 2024.2001.2029.577857. 10.1101/2024.01.29.577857.

10. DeWeirdt, P.C., Mahoney, E.M., and Laub, M.T. (2025). DefensePredictor: A Machine Learning Model to Discover Novel Prokaryotic Immune Systems. bioRxiv, 2025.2001.2008.631726. 10.1101/2025.01.08.631726.

11. Aravind, L., and Koonin, E.V. (1999). Novel Predicted RNA-Binding Domains Associated with the Translation Machinery. J Mol Evol 48, 291–302. 10.1007/PL00006472.

12. Ishitani, R., Nureki, O., Nameki, N., Okada, N., Nishimura, S., and Yokoyama, S. (2003). Alternative Tertiary Structure of tRNA for Recognition by a Posttranscriptional Modification Enzyme. Cell 113, 383–394. 10.1016/S0092-8674(03)00280-0.

13. Duan, J., Li, L., Lu, J., Wang, W., and Ye, K. (2009). Structural Mechanism of Substrate RNA Recruitment in H/ACA RNA-Guided Pseudouridine Synthase. Mol Cell 34, 427–439. 10.1016/j.molcel.2009.05.005.

14. Pérez-Arellano, I., Gallego, J., and Cervera, J. (2007). The PUA domain − a structural and functional overview. FEBS J 274, 4972–4984. 10.1111/j.1742-4658.2007.06031.x.

15. Bell, R.T., Wolf, Y.I., and Koonin, E.V. (2020). Modified base-binding EVE and DCD domains: striking diversity of genomic contexts in prokaryotes and predicted involvement in a variety of cellular processes. BMC Biol 18, 159. 10.1186/s12915-020-00885-2.

16. Hashimoto, H., Horton, J.R., Zhang, X., Bostick, M., Jacobsen, S.E., and Cheng, X. (2008). The SRA domain of UHRF1 flips 5-methylcytosine out of the DNA helix. Nature 455, 826–829. 10.1038/nature07280.

17. Iyer, L.M., Zhang, D., Maxwell Burroughs, A., and Aravind, L. (2013). Computational identification of novel biochemical systems involved in oxidation, glycosylation and other complex modifications of bases in DNA. Nucleic Acids Res 41, 7635–7655. 10.1093/nar/gkt573.

18. Lutz, T., Flodman, K., Copelas, A., Czapinska, H., Mabuchi, M., Fomenkov, A., He, X., Bochtler, M., and Xu, S.-y. (2019). A protein architecture guided screen for modification dependent restriction endonucleases. Nucleic Acids Res 47, 9761–9776. 10.1093/nar/gkz755.

19. Pabis, A., Duarte, F., and Kamerlin, S.C.L. (2016). Promiscuity in the Enzymatic Catalysis of Phosphate and Sulfate Transfer. Biochemistry 55, 3061–3081. 10.1021/acs.biochem.6b00297.

20. Zakataeva, N.P. (2021). Microbial 5′-nucleotidases: their characteristics, roles in cellular metabolism, and possible practical applications. Appl Microbiol Biotechnol 105, 7661–7681. 10.1007/s00253-021-11547-w.

21. Aravind, L., Iyer, L.M., and Burroughs, A.M. (2022). Discovering Biological Conflict Systems Through Genome Analysis: Evolutionary Principles and Biochemical Novelty. Ann Rev Biomed Data Sci 5, 367–391. 10.1146/annurev-biodatasci-122220-101119.

22. Zhang, J., Zhang, Y., and Inouye, M. (2003). Thermotoga maritima MazG Protein Has Both Nucleoside Triphosphate Pyrophosphohydrolase and Pyrophosphatase Activities. J Biol Chem 278, 21408–21414. 10.1074/jbc.M213294200.

23. Burroughs, A.M., Allen, K.N., Dunaway-Mariano, D., and Aravind, L. (2006). Evolutionary Genomics of the HAD Superfamily: Understanding the Structural Adaptations and Catalytic Diversity in a Superfamily of Phosphoesterases and Allied Enzymes. J Mol Biol 361, 1003–1034. 10.1016/j.jmb.2006.06.049.

24. Selengut, J.D. (2001). MDP-1 Is a New and Distinct Member of the Haloacid Dehalogenase Family of Aspartate-Dependent Phosphohydrolases. Biochemistry 40, 12704–12711. 10.1021/bi011405e.

25. Morais, M.C., Zhang, W., Baker, A.S., Zhang, G., Dunaway-Mariano, D., and Allen, K.N. (2000). The Crystal Structure of Bacillus cereus Phosphonoacetaldehyde Hydrolase: Insight into Catalysis of Phosphorus Bond Cleavage and Catalytic Diversification within the HAD Enzyme Superfamily. Biochemistry 39, 10385–10396. 10.1021/bi001171j.

26. Baker, A.S., Ciocci, M.J., Metcalf, W.W., Kim, J., Babbitt, P.C., Wanner, B.L., Martin, B.M., and Dunaway-Mariano, D. (1998). Insights into the Mechanism of Catalysis by the P−C Bond-Cleaving Enzyme Phosphonoacetaldehyde Hydrolase Derived from Gene Sequence Analysis and Mutagenesis. Biochemistry 37, 9305–9315. 10.1021/bi972677d.

27. Collet, J.-F., Gerin, I., Rider, M.H., Veiga-da-Cunha, M., and Van Schaftingen, E. (1997). Human l-3-phosphoserine phosphatase: sequence, expression and evidence for a phosphoenzyme intermediate. FEBS Lett 408, 281–284. 10.1016/S0014-5793(97)00438-9.

28. Hisano, T., Hata, Y., Fujii, T., Liu, J.-Q., Kurihara, T., Esaki, N., and Soda, K. (1996). Crystal Structure of L-2-Haloacid Dehalogenase from Pseudomonas sp. YL: An α/β hydrolase structure that is different from the α/β hydrolase fold. J Biol Chem 271, 20322–20330. 10.1074/jbc.271.34.20322.

29. Peisach, E., Selengut, J.D., Dunaway-Mariano, D., and Allen, K.N. (2004). X-ray Crystal Structure of the Hypothetical Phosphotyrosine Phosphatase MDP-1 of the Haloacid Dehalogenase Superfamily. Biochemistry 43, 12770–12779. 10.1021/bi0490688.

30. Koonin, E.V., and Tatusov, R.L. (1994). Computer Analysis of Bacterial Haloacid Dehalogenases Defines a Large Superfamily of Hydrolases with Diverse Specificity: Application of an Iterative Approach to Database Search. J Mol Biol 244, 125–132. 10.1006/jmbi.1994.1711.

31. Miller, E.S., Kutter, E., Mosig, G., Arisaka, F., Kunisawa, T., and Rüger, W. (2003). Bacteriophage T4 Genome. Microbiol Mol Biol Rev 67, 86–156. 10.1128/mmbr.67.1.86-156.2003.

32. Hercules, K., Munro, J.L., Mendelsohn, S., and Wiberg, J.S. (1971). Mutants in a Nonessential Gene of Bacteriophage T4 Which Are Defective in the Degradation of Escherichia coli Deoxyribonucleic Acid. J Virol 7, 95–105. 10.1128/jvi.7.1.95-105.1971.

33. Xu, C., Liu, K., Ahmed, H., Loppnau, P., Schapira, M., and Min, J. (2015). Structural Basis for the Discriminative Recognition of N^6^-Methyladenosine RNA by the Human YT521-B Homology Domain Family of Proteins. J Biol Chem 290, 24902–24913. 10.1074/jbc.M115.680389.

34. Bertonati, C., Punta, M., Fischer, M., Yachdav, G., Forouhar, F., Zhou, W., Kuzin, A.P., Seetharaman, J., Abashidze, M., Ramelot, T.A., et al. (2009). Structural genomics reveals EVE as a new ASCH/PUA-related domain. Prot Struct Funct Bioinf 75, 760–773. 10.1002/prot.22287.

35. Cui, Y., Dai, Z., Ouyang, Y., Fu, C., Wang, Y., Chen, X., Yang, K., Zheng, S., Wang, W., Tao, P., et al. (2025). Bacterial Hachiman complex executes DNA cleavage for antiphage defense. Nat Comm 16, 2604. 10.1038/s41467-025-57851-1.

36. Ho, C.-H., Wang, H.-C., Ko, T.-P., Chang, Y.-C., and Wang, A.H.J. (2014). The T4 Phage DNA Mimic Protein Arn Inhibits the DNA Binding Activity of the Bacterial Histone-like Protein H-NS. J Biol Chem 289, 27046–27054. 10.1074/jbc.M114.590851.

37. Wilkinson, M., Troman, L.A., Wan Nur Ismah, W.A.K., Chaban, Y., Avison, M.B., Dillingham, M.S., and Wigley, D.B. (2016). Structural basis for the inhibition of RecBCD by Gam and its synergistic antibacterial effect with quinolones. eLife 5, e22963. 10.7554/eLife.22963.

38. Walkinshaw, M.D., Taylor, P., Sturrock, S.S., Atanasiu, C., Berge, T., Henderson, R.M., Edwardson, J.M., and Dryden, D.T.F. (2002). Structure of Ocr from Bacteriophage T7, a Protein that Mimics B-Form DNA. Mol Cell 9, 187–194. 10.1016/S1097-2765(02)00435-5.

39. Chowdhury, S., Carter, J., Rollins, M.F., Golden, S.M., Jackson, R.N., Hoffmann, C., Nosaka, L.A., Bondy-Denomy, J., Maxwell, K.L., Davidson, A.R., et al. (2017). Structure Reveals Mechanisms of Viral Suppressors that Intercept a CRISPR RNA-Guided Surveillance Complex. Cell 169, 47–57.e11. 10.1016/j.cell.2017.03.012.

40. Taranenko, D., Kotovskaya, O., Kuznedelov, K., Yanovskaya, D., Demkina, A., Fardeeva, S., Mamontov, V., Vierra, K., Burman, N., Li, D., et al. (2025). A census of anti-CRISPR proteins reveals AcrIE9 as an inhibitor of Escherichia coli K12 Type IE CRISPR-Cas system. bioRxiv, 2025.2005.2007.652737. 10.1101/2025.05.07.652737.

41. Pinilla-Redondo, R., Shehreen, S., Marino, N.D., Fagerlund, R.D., Brown, C.M., Sørensen, S.J., Fineran, P.C., and Bondy-Denomy, J. (2020). Discovery of multiple anti-CRISPRs highlights anti-defense gene clustering in mobile genetic elements. Nat Comm 11, 5652. 10.1038/s41467-020-19415-3.

42. Mol, C., Arvai, A., Sanderson, R., Slupphaug, G., Kavli, B., Krokan, H., Mosbaugh, D., and Tainer, J. (1995). Crystal structure of human uracil-DNA glycosylase in complex with a protein inhibitor: protein mimicry of DNA. Cell 82, 701–708. 10.1016/0092-8674(95)90467-0.

43. Serrano-Heras, G., Ruiz-Masó, J.A., del Solar, G., Espinosa, M., Bravo, A., and Salas, M. (2007). Protein p56 from the Bacillus subtilis phage LJ29 inhibits DNA-binding ability of uracil-DNA glycosylase. Nucleic Acids Res 35, 5393–5401. 10.1093/nar/gkm584.

44. Wang, Z., Wang, H., Mulvenna, N., Sanz-Hernandez, M., Zhang, P., Li, Y., Ma, J., Wang, Y., Matthews, S., Wigneshweraraj, S., and Liu, B. (2021). A Bacteriophage DNA Mimic Protein Employs a Non-specific Strategy to Inhibit the Bacterial RNA Polymerase. Front Microbiol 12, 2021. 10.3389/fmicb.2021.692512.

45. McMahon, S.A., Roberts, G.A., Johnson, K.A., Cooper, L.P., Liu, H., White, J.H., Carter, L.G., Sanghvi, B., Oke, M., Walkinshaw, M.D., et al. (2009). Extensive DNA mimicry by the ArdA anti-restriction protein and its role in the spread of antibiotic resistance. Nucleic Acids Res 37, 4887–4897. 10.1093/nar/gkp478.

46. Kang, H., Gao, A., and Zhu, Y. (2025). Bacterial restriction-modification systems: mechanisms of defense against phage infection. Biophys Rep 11, 330–343. 10.52601/bpr.2025.240070.

47. Niault, T., van Houte, S., Westra, E., and Swarts, D.C. (2025). Evolution and ecology of anti-defence systems in phages and plasmids. Curr Biol 35, R32–R44. 10.1016/j.cub.2024.11.033.

48. Wang, Y., Wang, C., Guan, Z., Cao, J., Xu, J., Wang, S., Cui, Y., Wang, Q., Chen, Y., Yin, Y., et al. (2024). DNA methylation activates retron Ec86 filaments for antiphage defense. Cell Rep 43. 10.1016/j.celrep.2024.114857.

49. Stella, G., Ye, L., Brady, S.F., and Marraffini, L. (2025). CARF-HAD phosphatase effectors provide immunity during the type III-A CRISPR–Cas response. Nucleic Acids Res 53. 10.1093/nar/gkaf1363.

50. Zhou, Y.J., Yang, W., Wang, L., Zhu, Z., Zhang, S., and Zhao, Z.K. (2013). Engineering NAD+ availability for Escherichia coli whole-cell biocatalysis: a case study for dihydroxyacetone production. Microb Cell Fact 12, 103. 10.1186/1475-2859-12-103.

51. Rodionov, D.A., Li, X., Rodionova, I.A., Yang, C., Sorci, L., Dervyn, E., Martynowski, D., Zhang, H., Gelfand, M.S., and Osterman, A.L. (2008). Transcriptional regulation of NAD metabolism in bacteria: genomic reconstruction of NiaR (YrxA) regulon. Nucleic Acids Res 36, 2032–2046. 10.1093/nar/gkn046.

52. Schaaper, R.M., and Mathews, C.K. (2013). Mutational consequences of dNTP pool imbalances in E. coli. DNA Repair 12, 73–79. 10.1016/j.dnarep.2012.10.011.

53. Poli, J., Tsaponina, O., Crabbé, L., Keszthelyi, A., Pantesco, V., Chabes, A., Lengronne, A., and Pasero, P. (2012). dNTP pools determine fork progression and origin usage under replication stress. EMBO J 31, 883–894. 10.1038/emboj.2011.470.

54. Xu, H., He, X., Zheng, H., Huang, L.J., Hou, F., Yu, Z., de la Cruz, M.J., Borkowski, B., Zhang, X., Chen, Z.J., and Jiang, Q.-X. (2014). Structural basis for the prion-like MAVS filaments in antiviral innate immunity. eLife 3, e01489. 10.7554/eLife.01489.

55. Liu, S., Yang, B., Hou, Y., Cui, K., Yang, X., Li, X., Chen, L., Liu, S., Zhang, Z., Jia, Y., et al. (2023). The mechanism of STING autoinhibition and activation. Mol Cell 83, 1502–1518.e1510. 10.1016/j.molcel.2023.03.029.

56. Morehouse, B.R., Yip, M.C.J., Keszei, A.F.A., McNamara-Bordewick, N.K., Shao, S., and Kranzusch, P.J. (2022). Cryo-EM structure of an active bacterial TIR–STING filament complex. Nature 608, 803–807. 10.1038/s41586-022-04999-1.

57. Hogrel, G., Guild, A., Graham, S., Rickman, H., Grüschow, S., Bertrand, Q., Spagnolo, L., and White, M.F. (2022). Cyclic nucleotide-induced helical structure activates a TIR immune effector. Nature 608, 808–812. 10.1038/s41586-022-05070-9.

58. Baca, C.F., Majumder, P., Hickling, J.H., Patel, D.J., and Marraffini, L.A. (2025). Cat1 forms filament networks to degrade NAD^+^ during the type III CRISPR-Cas antiviral response. Science 388, eadv9045. 10.1126/science.adv9045.

59. Wang, F., Xu, H., Zhang, C., Xue, J., and Li, Z. (2025). Target DNA-induced filament formation and nuclease activation of SPARDA complex. Cell Res 35, 510–519. 10.1038/s41422-025-01100-z.

60. Kanevskaya, A., Narwal, M., Lisitskaya, L., Moiseenko, A.V., Sokolova, O.S., Sluchanko, N.N., Murakami, K.S., and Kulbachinskiy, A. (2025). Argonaute-HNH filaments triggered by invader DNA confer bacterial immunity. Nat Comm. 10.1038/s41467-025-66189-7.

61. Wang, Y., Tian, Y., Yang, X., Yu, F., and Zheng, J. (2025). Filamentation activates bacterial Avs5 antiviral protein. Nat Comm 16, 2408. 10.1038/s41467-025-57732-7.

62. Cheng, R., Huang, F., Lu, X., Yan, Y., Yu, B., Wang, X., and Zhu, B. (2023). Prokaryotic Gabija complex senses and executes nucleotide depletion and DNA cleavage for antiviral defense. Cell Host Microbe 31, 1331–1344.e1335. 10.1016/j.chom.2023.06.014.

63. Aframian, N., Omer Bendori, S., Hen, T., Guler, P., and Eldar, A. (2025). Expression level of anti-phage defence systems controls a trade-off between protection range and autoimmunity. Nat Microbiol 10, 1954–1962. 10.1038/s41564-025-02063-y.

64. Osterman, I., Hurieva, B., Moses, S., Falkovich, A.H., Itkin, M., Malitsky, S., Yirmiya, E., and Sorek, R. (2025). Bacteria sense virus-induced genome degradation via methylated mononucleotides. bioRxiv, 2025.2011.2005.686725. 10.1101/2025.11.05.686725.

65. Isaev, A., Drobiazko, A., Sierro, N., Gordeeva, J., Yosef, I., Qimron, U., Ivanov, N.V., and Severinov, K. (2020). Phage T7 DNA mimic protein Ocr is a potent inhibitor of BREX defence. Nucleic Acids Res 48, 5397–5406. 10.1093/nar/gkaa290.

66. Bandyopadhyay, P.K., Studier, F.W., Hamilton, D.L., and Yuan, R. (1985). Inhibition of the type I restriction-modification enzymes EcoB and EcoK by the gene 0.3 protein of bacteriophage T7. J Mol Biol 182, 567–578. 10.1016/0022-2836(85)90242-6.

67. Altschul, S.F., Madden, T.L., Schäffer, A.A., Zhang, J., Zhang, Z., Miller, W., and Lipman, D.J. (1997). Gapped BLAST and PSI-BLAST: a new generation of protein database search programs. Nucleic Acids Res 25, 3389–3402. 10.1093/nar/25.17.3389.

68. Johnson, L.S., Eddy, S.R., and Portugaly, E. (2010). Hidden Markov model speed heuristic and iterative HMM search procedure. BMC Bioinf 11, 431. 10.1186/1471-2105-11-431.

69. Steinegger, M., and Söding, J. (2017). MMseqs2 enables sensitive protein sequence searching for the analysis of massive data sets. Nat Biotechnol 35, 1026–1028. 10.1038/nbt.3988.

70. Katoh, K., Misawa, K., Kuma, K.i., and Miyata, T. (2002). MAFFT: a novel method for rapid multiple sequence alignment based on fast Fourier transform. Nucleic Acids Res 30, 3059–3066. 10.1093/nar/gkf436.

71. Sonnhammer, E.L.L., Eddy, S.R., and Durbin, R. (1997). Pfam: A comprehensive database of protein domain families based on seed alignments. Prot Struct Funct Bioinf 28, 405–420. 10.1002/(SICI)1097-0134(199707)28:3<405::AID-PROT10>3.0.CO;2-L.

72. Söding, J., Biegert, A., and Lupas, A.N. (2005). The HHpred interactive server for protein homology detection and structure prediction. Nucleic Acids Res 33, W244–W248. 10.1093/nar/gki408.

73. Steinegger, M., Meier, M., Mirdita, M., Vöhringer, H., Haunsberger, S.J., and Söding, J. (2019). HH-suite3 for fast remote homology detection and deep protein annotation. BMC Bioinformatics 20, 473. 10.1186/s12859-019-3019-7.

74. Blondel, V.D., Guillaume, J.-L., Lambiotte, R., and Lefebvre, E. (2008). Fast unfolding of communities in large networks. J Stat Mech: Theory Exp 2008, P10008. 10.1088/1742-5468/2008/10/P10008.

75. Fruchterman, T.M.J., and Reingold, E.M. (1991). Graph drawing by force-directed placement. Softw: Pract Exp 21, 1129–1164. 10.1002/spe.4380211102.

76. Kamada, T., and Kawai, S. (1989). An algorithm for drawing general undirected graphs. Inform Proc Lett 31, 7–15. 10.1016/0020-0190(89)90102-6.

77. Antonov, M., Csárdi, G., Horvát, S., Müller, K., Nepusz, T., Noom, D., Salmon, M., Traag, V., Welles, B., and F, Z. (2023). igraph enables fast and robust network analysis across programming languages. arXiv 2311.*10260*. 10.48550/arXiv.2311.10260.

78. Hagberg, A.A., Schult, D.A., and Swart, P.J. (2008). Exploring network structure, dynamics, and function using NetworkX. held in Pasadena, CA USA, G. Varoquaux, T. Vaught, and J. Millman, eds. pp. 11–15.

79. Prjibelski, A., Antipov, D., Meleshko, D., Lapidus, A., and Korobeynikov, A. (2020). Using SPAdes De Novo Assembler. Curr Prot Bioinf 70, e102. 10.1002/cpbi.102.

80. Li, H. (2013). Aligning sequence reads, clone sequences and assembly contigs with BWA-MEM. arXiv: Genomics.

81. Danecek, P., Bonfield, J.K., Liddle, J., Marshall, J., Ohan, V., Pollard, M.O., Whitwham, A., Keane, T., McCarthy, S.A., Davies, R.M., and Li, H. (2021). Twelve years of SAMtools and BCFtools. GigaScience 10. 10.1093/gigascience/giab008.

82. van der Auwera, G., and O’Connor, B. (2020). Genomics in the Cloud: Using Docker, GATK, and WDL in Terra (1st Edition) (O’Reilly Media).

83. Punjani, A., Rubinstein, J.L., Fleet, D.J., and Brubaker, M.A. (2017). cryoSPARC: algorithms for rapid unsupervised cryo-EM structure determination. Nat Methods 14, 290–296. 10.1038/nmeth.4169.

84. Zivanov, J., Nakane, T., Forsberg, B.O., Kimanius, D., Hagen, W.J., Lindahl, E., and Scheres, S.H. (2018). New tools for automated high-resolution cryo-EM structure determination in RELION-3. Elife 7. 10.7554/eLife.42166.

85. Abramson, J., Adler, J., Dunger, J., Evans, R., Green, T., Pritzel, A., Ronneberger, O., Willmore, L., Ballard, A.J., Bambrick, J., et al. (2024). Accurate structure prediction of biomolecular interactions with AlphaFold 3. Nature 630, 493–500. 10.1038/s41586-024-07487-w.

86. Emsley, P., Lohkamp, B., Scott, W.G., and Cowtan, K. (2010). Features and development of Coot. Acta Crystallogr D Biol Crystallogr 66, 486–501. 10.1107/S0907444910007493.

87. Adams, P.D., Afonine, P.V., Bunkoczi, G., Chen, V.B., Davis, I.W., Echols, N., Headd, J.J., Hung, L.W., Kapral, G.J., Grosse-Kunstleve, R.W., et al. (2010). PHENIX: a comprehensive Python-based system for macromolecular structure solution. Acta Crystallogr D Biol Crystallogr 66, 213–221. 10.1107/S0907444909052925.

88. Pettersen, E.F., Goddard, T.D., Huang, C.C., Meng, E.C., Couch, G.S., Croll, T.I., Morris, J.H., and Ferrin, T.E. (2021). UCSF ChimeraX: Structure visualization for researchers, educators, and developers. Protein Sci 30, 70–82. 10.1002/pro.3943.

89. Pettersen, E.F., Goddard, T.D., Huang, C.C., Couch, G.S., Greenblatt, D.M., Meng, E.C., and Ferrin, T.E. (2004). UCSF Chimera--a visualization system for exploratory research and analysis. J Comput Chem 25, 1605–1612. 10.1002/jcc.20084.

