## Supplementary material for "A methylome-derived m^6^-dAMP trigger assembles a PUA-Cal-HAD immune filament that depletes dNTPs to abort phage infection": Figure S

### Contents

**Figure S1** Diverse genomic architectures and sequence conservation of PUA-like Calcineurin-CE defence loci, Related to Figure 1

**Figure S2** The anti-phage activity of PUA-Cal-HAD, Related to Figure 2

**Figure S3** Oligomeric behaviour and cryo-EM data processing of PUA-Cal, Related to Figure 3

**Figure S4** Interfaces and conformational asymmetry within the PUA-Cal dimer, Related to Figure 3

**Figure S5** Ligand binding and catalytic architecture of the Calcineurin-CE domain and structural features and interaction properties of the HAD phosphatase, Related to Figures 3 and 4

**Figure S6** *In vivo* and *in vitro* activity of PUA-Cal-HAD on different nucleotide-derived substrates, Related to Figure 5

**Figure S7** $m^6$-dAMP-dependent fibre formation and interfacial interactions in the PUA-Cal system, Related to Figure 6

**Table S1** Genomic coordinates of predicted anti-phage systems featuring a PUA-like domain, Related to Figure 1 (*provided separately*)

**Table S2** LC-MS metabolic profiling of bacterial cultures containing PUA-Cal-HAD, Related to Figure 5

**Table S3** T2 escape mutants of PUA-Cal-HAD

**Table S4** List of strains and phages used in this study, Related to STAR Methods

**Table S5** List of plasmids used in this study, Related to STAR Methods

**Table S6** List of primers used in this study, Related to STAR Methods (*provided separately)*

**Table S7** Cryo-EM data collection and refinement statistics, Related to STAR Methods (*provided separately*)

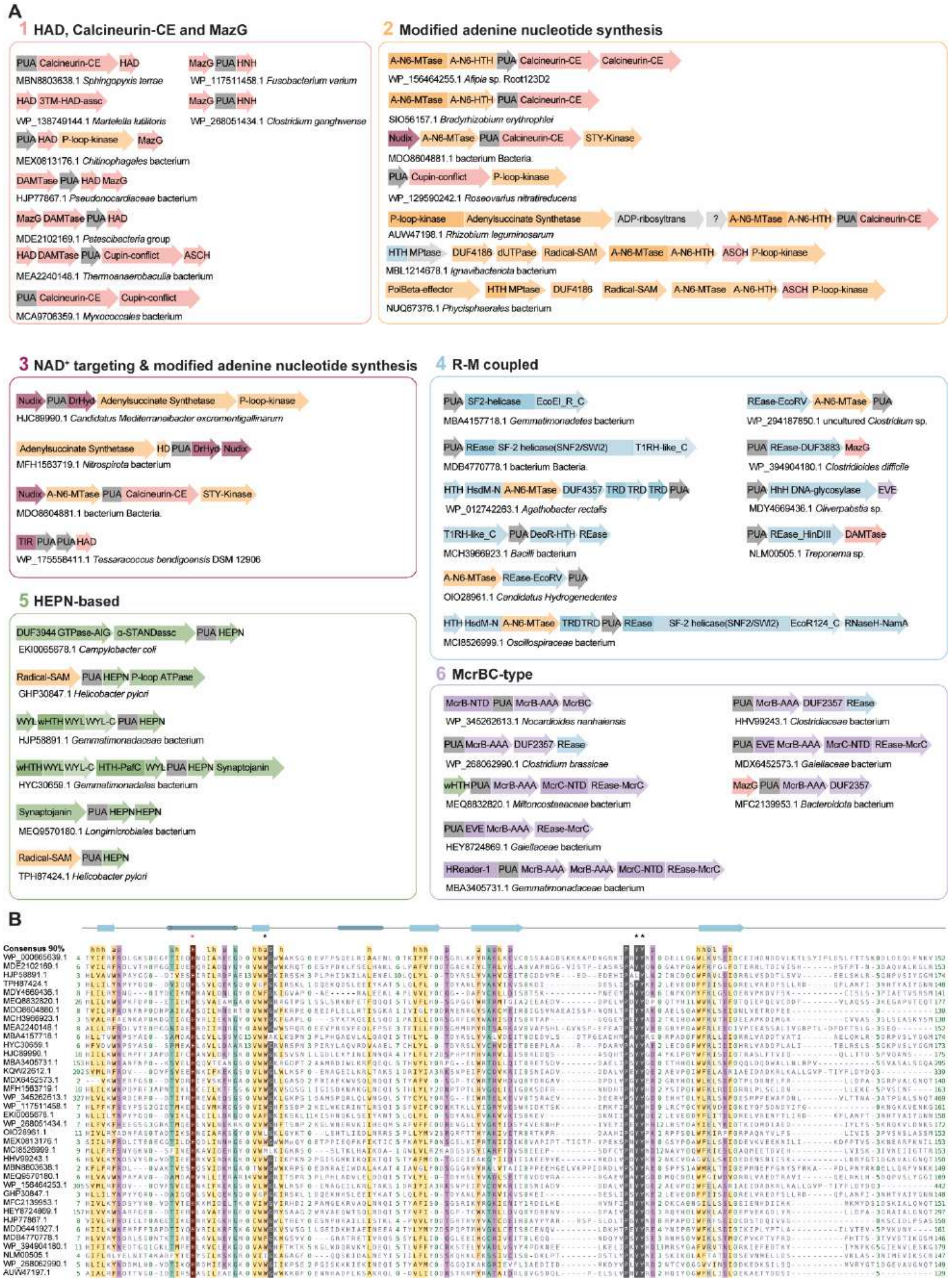

**Figure S1.** Diverse genomic architectures and sequence conservation of PUA-like Calcineurin-CE defence loci, Related to Figure 1

**(A)** Representative genomic contexts for the PUA-like domain family under consideration, grouped by shared functional themes. Genes are depicted as box arrows, with arrowheads indicating the 3′ ends of genes. Genes encoding multi-domain proteins are segmented into labelled regions corresponding to individual domains. Domains associated with modified nucleotide synthesis are highlighted in orange, whereas domains associated with $NAD^+$ targeting are highlighted in magenta.

**(B)** Multiple alignment of the PUA-like domain family considered in this study. The asterisks above the residues mark the positions predicted to be involved in recognition of modified nucleobase. The alignment is coloured according to the 90% consensus.

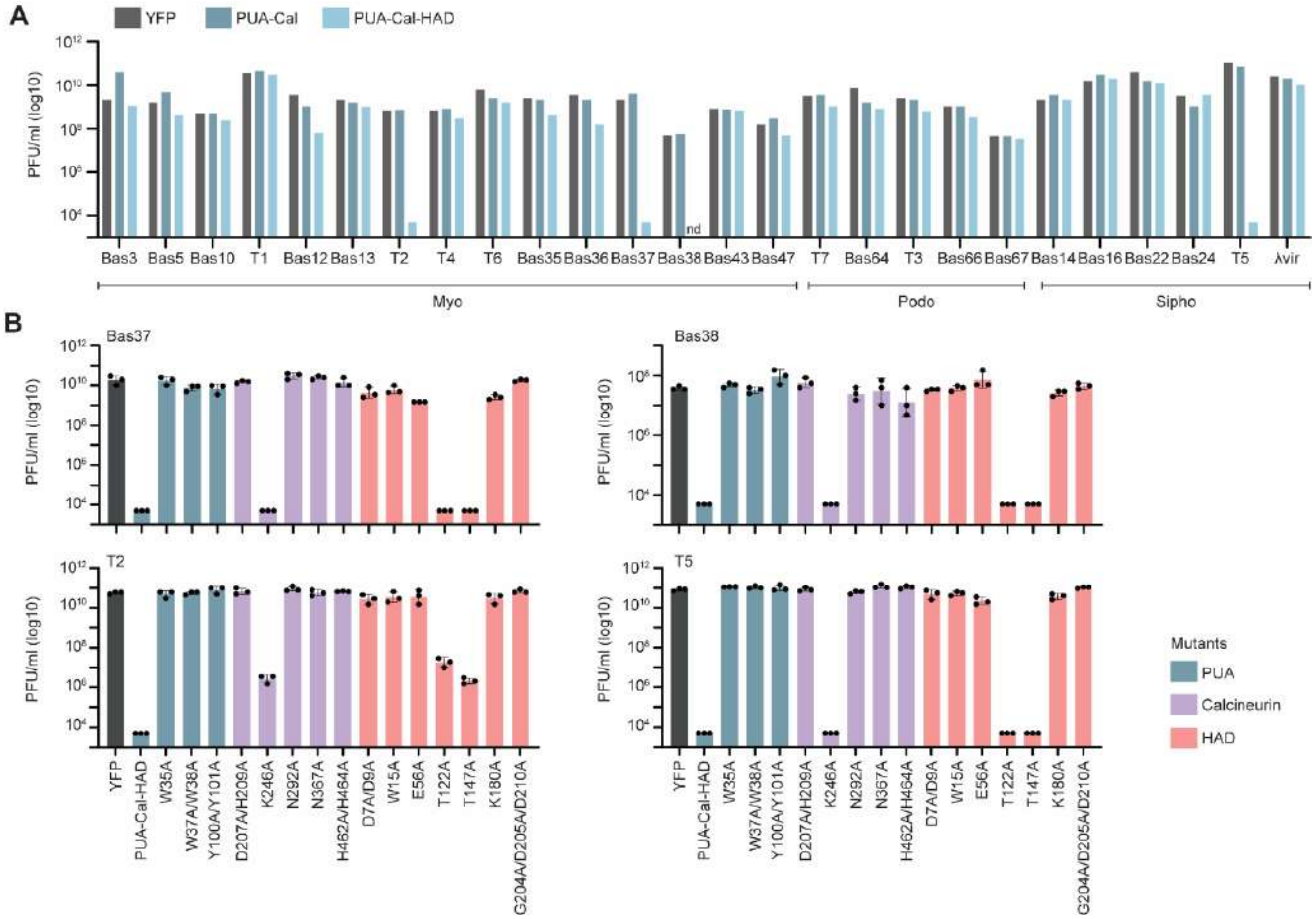


**Figure S2** The anti-phage activity of PUA-Cal-HAD, Related to Figure 2

**(A)** Infectivity (PFU/ml) of a panel of phages against YFP- (control), PUA-Cal-, and PUA-Cal-HAD-expressing cells. nd, plaques not detected.

**(B)** Infectivity (PFU/ml) of phages Bas37, Bas38, T2 and T5 against PUA-Cal-HAD mutants.

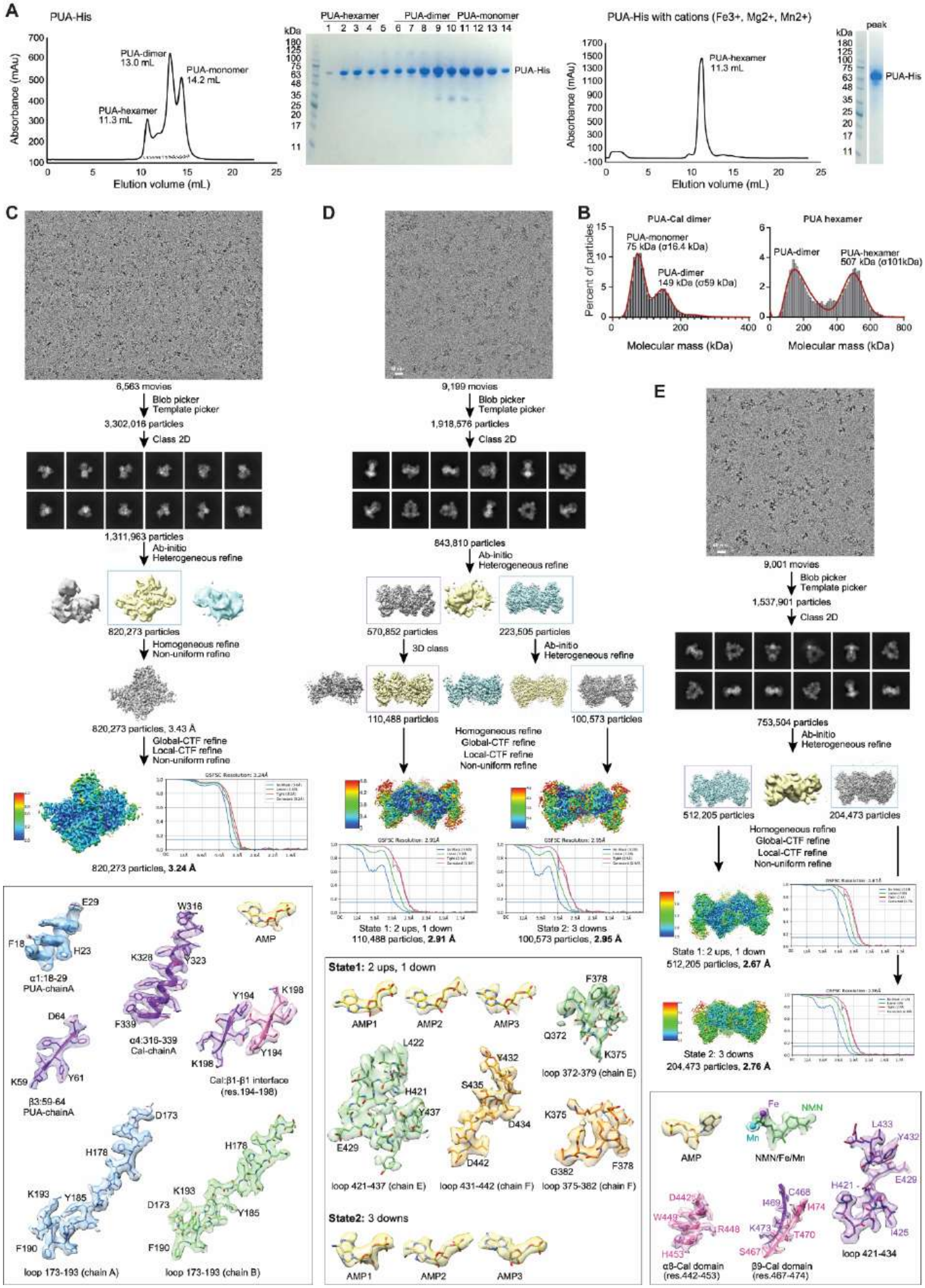

**Figure S3** Oligomeric behaviour and cryo-EM data processing of PUA-Cal, Related to Figure 3

**(A)** Size-exclusion chromatography (SEC) analysis of PUA-His. Left, analysis of PUA-His on a Superdex 200 Increase column reveals three species corresponding to hexameric, dimeric, and monomeric assemblies. SDS-PAGE analysis of peak fractions confirms the presence of PUA-His in all oligomeric states. Right, analysis of PUA-His in the presence of divalent metal ions shows a predominant hexameric species.

**(B)** Mass photometry analysis of PUA-Cal dimer and hexameric fractions shown as percent of particles per molecular mass in kDa.

**(C)** Cryo-EM data processing and representative densities of the AMP-bound PUA-Cal dimer. Top, representative cryo-EM micrograph. Middle, cryo-EM image processing workflow, yielding a final reconstruction at 3.24 Å resolution. Bottom, representative local cryo-EM densities highlighting well-resolved important regions, including the AMP-binding site, loop 173-193 in both protomers, and selected secondary structure elements in the PUA and Cal domains.

**(D)** Cryo-EM data processing and representative densities of the AMP-bound PUA-Cal hexamer. Top, representative cryo-EM micrograph. Middle, cryo-EM image processing workflow, yielding two predominant conformational states of the hexamer at overall resolutions of 2.91 Å (State 1: 2 ups, 1 down) and 2.95 Å (State 2: 3 downs). Bottom, representative local cryo-EM densities, highlighting well-resolved important features, including AMP molecules bound at the PUA domains, conformational differences in loops 375-382, 372-379, 421-437, and 431-442, and secondary-structure elements at Cal-Cal interfaces.

**(E)** Cryo-EM data processing and representative densities of the ion-bound PUA-Cal hexamer. Top, representative cryo-EM micrograph. Middle, cryo-EM image processing workflow, resulting in two major conformational states refined to overall resolutions of 2.67 Å (State 1: 2 ups, 1 down) and 2.76 Å (State 2: 3 downs). Bottom, representative local cryo-EM densities illustrating well-resolved ligands and protein regions, including NMN coordinated with Fe/Mn in the Cal active site, AMP bound at the PUA domain, and flexible Cal loops (residues 421-434, 442-453, and 467-474).

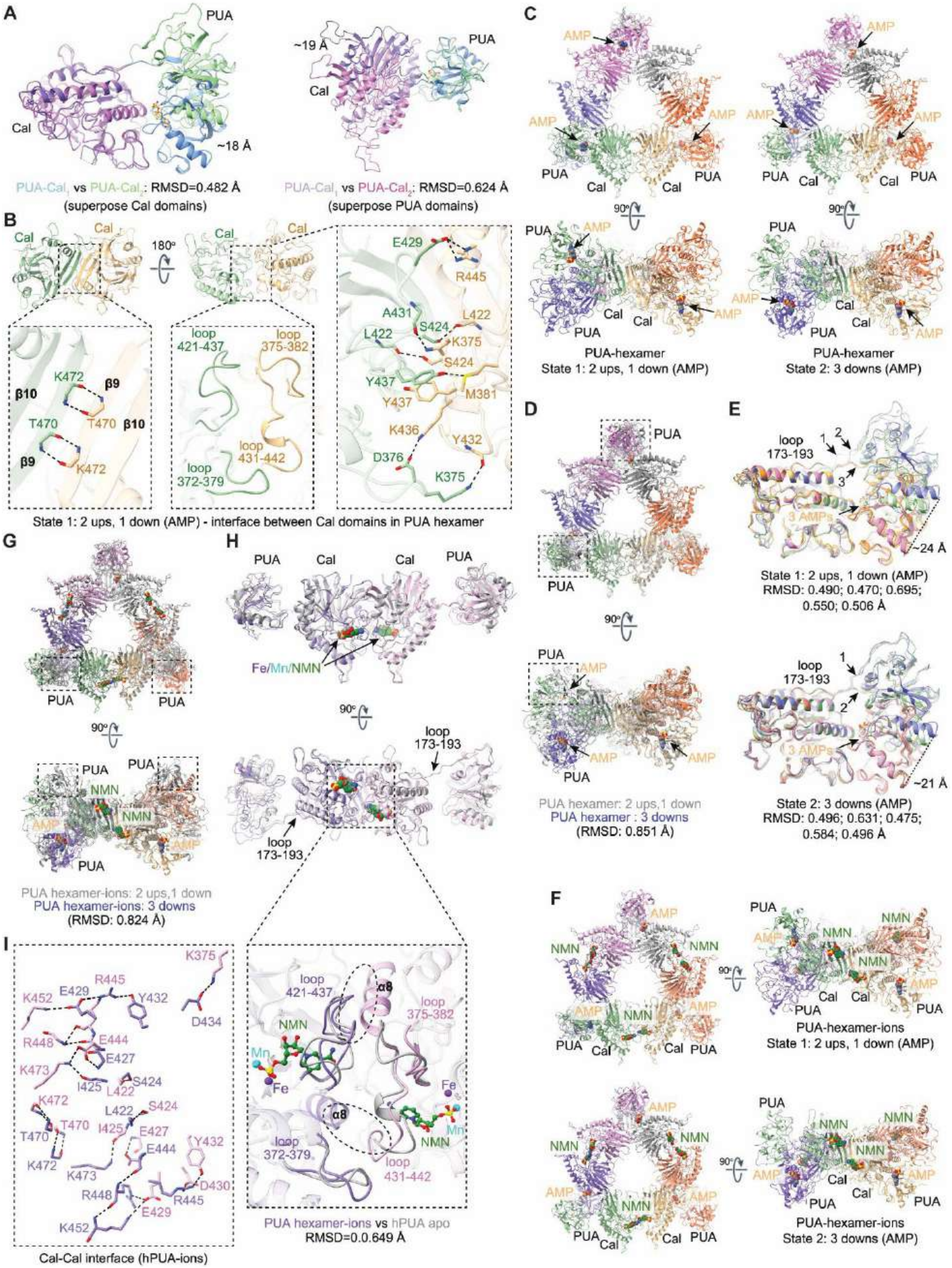


**Figure S4** Interfaces and conformational asymmetry within the PUA-Cal dimer, Related to Figure 3

**(A)** Structural superposition of the two PUA-Cal promoters within the dimer, based on the Cal domains (left) or PUA domains (right).

**(B)** Close-up views of the Cal-Cal interfaces within the AMP-bound hexamer (State 1). Insets highlight secondary-structure interactions involving β9-β10 main-chain contacts (left) and loop-loop interactions between loop 421-437 of one promoter and loop 372-379 of the adjacent protomer (middle), with detailed interacting residues shown in the right panel.

**(C)** Cryo-EM reconstructions of the AMP-bound PUA-Cal hexamer reveal two predominant conformational states. Left, State 1, in which two PUA domains adopt an up conformation and bind AMP, while one PUA domain adopts a down conformation within each dimeric PUA of the hexamer. Right, State 2, in which all three PUA domains adopt the down conformation and bind AMP. Structures are shown in two orientations, with AMP bound at the PUA domains.

**(D)** Superposition of the two AMP-bound hexamer states. Dashed boxes highlight PUA domains that undergo conformational switching between up and down positions.

**(E)** Structural comparison of PUA domains from the two AMP-bound hexamer states by superposition of the Cal domains. The comparison highlights movements of loop 173-193 at the PUA-Cal junction, together with adjacent secondary-structure elements, associated with transitions between up and down conformations. Distances between corresponding PUA domains are indicated.

**(F)** Cryo-EM reconstructions of the ion-bound PUA-Cal hexamer in two conformational states. Top, State 1, in which two PUA domains adopt an up conformation, and one adopts a down conformation within each trimeric half of the hexamer. Bottom, State 2, in which all three PUA domains adopt the down conformation. AMP is bound at the PUA domains and NMN is bound at the Cal active sites. Structures are shown in two orientations.

**(G)** Superposition of the two ion-bound hexamer states. Dashed boxes highlight PUA domains that undergo conformational switching between up and down positions.

**(H)** Structural comparison of the ion-bound hexamer and the apo PUA-Cal hexamer by superposition of the Cal domains. The PUA-Cal junction loop (residues 173-195) remains largely conserved between states, while binding of metal ions and NMN induces local rearrangements in loop 431-442 and helix α8, which move closer and form new contacts.

**(I)** Close-up view of the Cal-Cal interface in the ion-bound hexamer. Key interacting residues are sown, with the State 1 configuration used as a representative example.

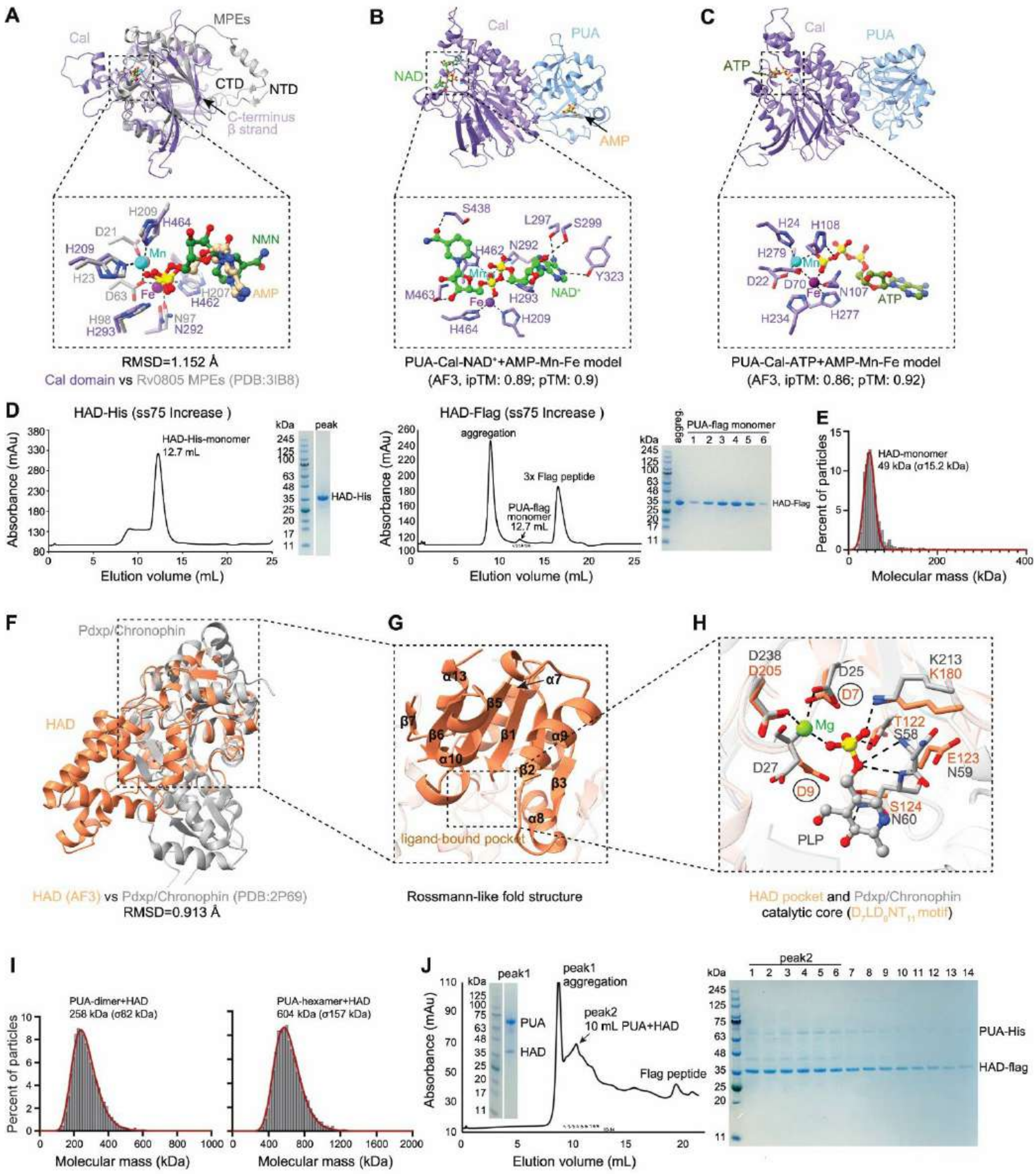


**Figure S5** Ligand binding and catalytic architecture of the Calcineurin-CE domain and structural features and interaction properties of the HAD phosphatase, Related to Figures 3 and 4

**(A)** Structural superposition of the Cal domain with a representative metal-dependent phosphoesterase (MPE) from *Mycobacterium tuberculosis* (Rv0805, PDB: 3IB8) showing strong overall similarity and supporting assignment of the Cal domain to the MPE/Calcineurin-CE family. N-terminal (NTD) and C-terminal (CTD) regions are indicated. At the bottom, comparison of the active sites of Rv0805 and the Cal domain. Conserved metal-coordinating residues and catalytic motifs are shown, highlighting the shared binuclear metal-centre architecture.

**(B)** AlphaFold 3 (AF3) model of the PUA-Cal fusion protein with $NAD^+$ bound in the Cal active site and AMP bound in the PUA domain. At the bottom, a close-up view of the predicted $NAD^+$-binding pocket in the Cal domain.

**(C)** AF3 model of the PUA-Cal protein with ATP bound in the Cal active site and AMP bound in the PUA domain. At the bottom, a close-up view of the ATP-binding pocket in the Cal domain.

**(D)** Size-exclusion chromatography analysis of HAD-His (left) and HAD-Flag (right) on a Superdex 75 Increase column. SDS-PAGE analysis of peak fractions confirming the identity of HAD-His and HAD-Flag.

**(E)** Mass photometry analysis HAD, shown as percent of particles per molecular mass in kDa.

**(F)** Structural superposition of the HAD domain with the human Pdxp/Chronophin phosphatase (PDB: 2P69) of the HAD superfamily.

**(G)** Close-up view of the HAD core reveals a Rossmann-like fold characteristic of HAD-family phosphatases.

**(H)** Detailed view of the HAD catalytic pocket, highlighting the conserved $D_7LD_9NT_{11}$ motif and surrounding residues that coordinate the $Mg^{2+}$ ion and support phosphatase activity. Key catalytic residues within the motif (D7 and D9) are circled.

**(I)** Mass photometry analysis PUA-Cal dimeric and hexameric fractions, together with HAD, shown as percent of particles per molecular mass in kDa.

**(J)** Size-exclusion chromatography analysis of co-expressed PUA-His and HAD-Flag proteins. SDS-PAGE analysis of peak fractions confirms the presence of both PUA-His and HAD-Flag, suggesting an interaction between the two proteins in solution, with a propensity to aggregate.

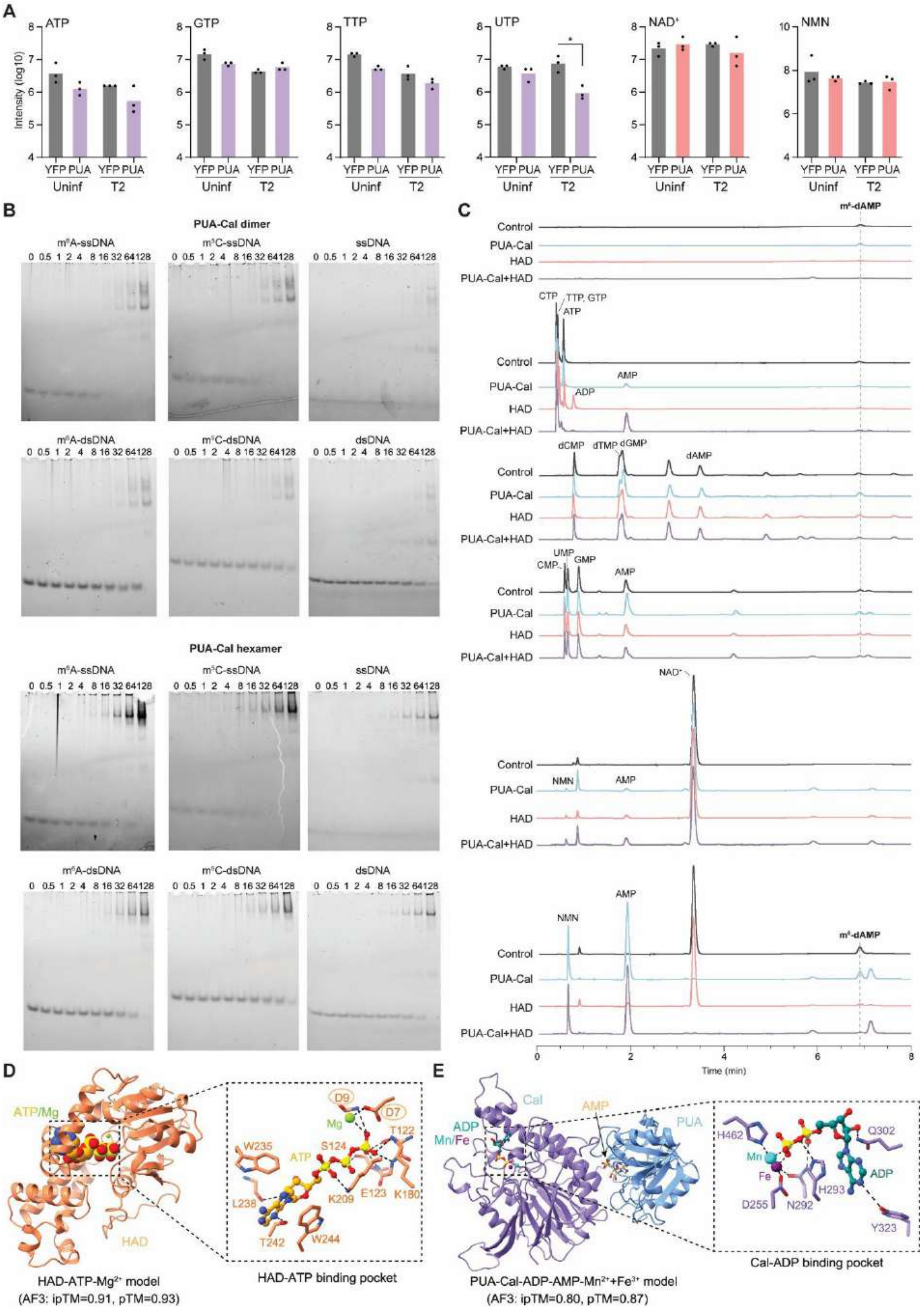


**Figure S6** *In vivo* and *in vitro* activity of PUA-Cal-HAD on different nucleotide-derived substrates,

Related to Figure 5

**(A)** Quantification of nucleotide and nucleotide-derived metabolites by LC-MS, shown as relative intensity levels in log10, in cells expressing YFP or PUA-Cal-HAD upon infection with phage T2.

**(B)** Electrophoretic mobility shift assay (EMSA) of PUA-Cal dimer and hexamer with different unmethylated and methylated DNA substrates.

**(C)** LC-MS measurement of PUA-Cal, HAD, or PUA-Cal-HAD *in vitro* activity on different nucleotide substrates in the presence of activator $m^6$-dAMP.

**(D)** AlphaFold 3 (AF3) model of HAD bound to ATP-$Mg^{2+}$, highlighting the conserved HAD catalytic pocket. Residues surrounding the DxDx(T/N) catalytic motif are shown as sticks, with the invariant D7 and D9 residues circled.

**(E)** AF3 model of the PUA-Cal complex bound to ADP and AMP. ADP coordinated by the Cal binuclear metal centre is shown as cyan sticks, AMP bound within the PUA pocket is shown as wheat-coloured sticks, and metal ions are shown as dark purple and cyan spheres.

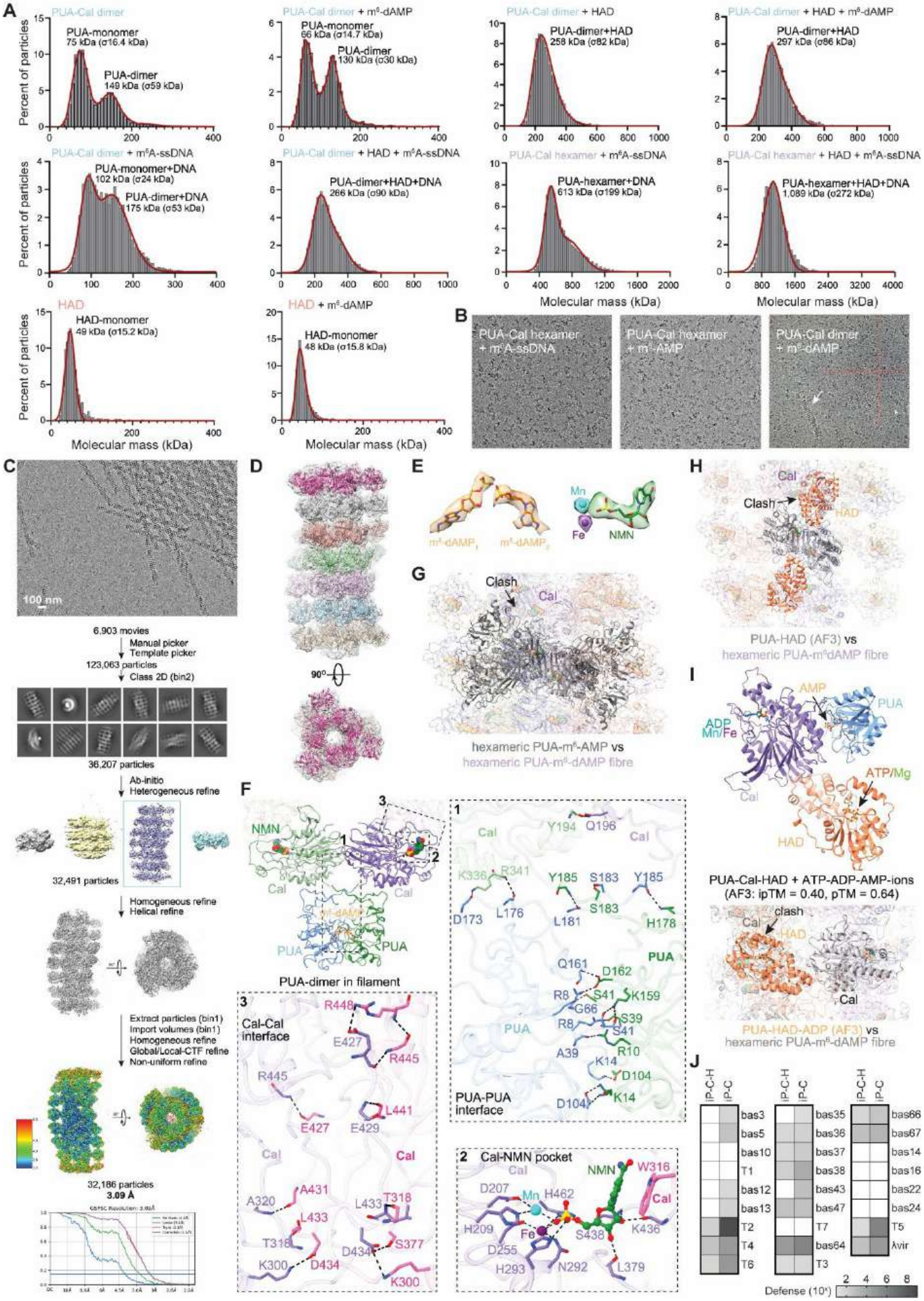


**Figure S7** m6-dAMP-dependent fibre formation and interfacial interactions in the PUA-Cal system,

Related to Figure 6

**(A)** Mass photometry analysis of PUA-Cal, HAD, or PUA-Cal and HAD in the presence and absence of $m^6$-dAMP and $m^6$A-containing single stranded DNA ($m^6$A-ssDNA). Results are shown as percent of particles per molecular mass in kDa.

**(B)** Representative cryo-EM raw micrographs of PUA-Cal assemblies in the presence of $m^6$A-ssDNA, $m^6$-AMP, or $m^6$-dAMP.

**(C)** Cryo-EM data processing and reconstruction of the $m^6$-dAMP-induced PUA-Cal filament. Top, representative micrograph. Bottom, cryo-EM image processing workflow for filament reconstruction, resulting in a final filament reconstruction at 3.04 Å resolution.

**(D)** Final cryo-EM reconstruction of the PUA-Cal filament shown in side and top views, with individual hexameric layers coloured separately to illustrate the helical organisation.

**(E)** Density fitting of ligand geometries in the filament state. Left, cryo-EM density corresponding to the two $m^6$-dAMP molecules bound within a PUA dimer in the filament assembly, illustrating their symmetric arrangement and close proximity. Right, cryo-EM density of NMN bound within the Cal active site together with the binuclear metal centre.

**(F)** Interface organisation within the $m^6$-dAMP-induced filament, with the two promoters shown in distinct colours. Insets highlight interaction networks that stabilise the filament state. (1) Expanded PUA-PUA interface, showing an extensive network of hydrogen bonds and salt bridges formed between residues from the two opposing PUA promoters. (2) Cal-NMN active site, illustrating conserved coordination of NMN by the binuclear metal centre within the Cal domain. (3) Cal-Cal interface, highlighting conserved polar interactions and salt bridges between neighbouring Cal promoters that stabilise inter-hexamer while preserving the catalytic core architecture.

**(G)** Structural superposition of the $m^6$-AMP bound PUA-Cal hexamer with the $m^6$-dAMP-induced filament model. Steric clashes (black arrow) are observed between the PUA domain from the hexamer and a neighbouring Cal domain from the filament, indicating that the $m^6$-AMP bound hexamer conformation is incompatible with filament assembly, whereas the $m^6$-dAMP bound state supports higher-order polymerisation.

**(H)** Superposition of the AlphaFold 3 (AF3) predicted PUA-HAD complex onto the cryo-EM $m^6$-dAMP-bound PUA-Cal filament reveals steric clashes, indicating that the predicted PUA-HAD arrangement is incompatible with the filament architecture and suggesting that HAD association is dynamic and repositioned during filament formation.

**(I)** AF3 model of the PUA-Cal-HAD assembly with ATP, ADP, and AMP bound. Superposition of this model onto the $m^6$-dAMP-induced filament reveals extensive steric clashes, indicating that HAD binding would require conformational rearrangement to be compatible with the filament state.

**(J)** Ten-fold defence of inducible PUA-Cal and PUA-Cal-HAD from ECOR8 against a panel of distinct phages.

**Table S2** LC-MS metabolic profiling of bacterial cultures containing PUA-Cal-HAD, Related to Figure 5

| | Relative intensity | | | | | | | | | |
|---|---|---|---|---|---|---|---|---|---|---|
| | **dATP** | **dGTP** | **dCTP** | **TTP** | **ATP** | **GTP** | **CTP** | **UTP** | **NAD+** | **NMN** |
| **YFP** | 2.2E7 | 1.5E7 | 3.4E6 | 1.3E7 | 8.7E6 | 2.2E7 | nd | 5.3E6 | 2.8E7 | 3.9E7 |
| | 2.0E7 | 2.5E7 | 4.5E6 | 1.5E7 | 2.0E6 | 9.9E6 | nd | 6.1E6 | 2.4E7 | 4.8E8 |
| | 2.3E7 | 1.4E7 | 3.2E6 | 1.5E7 | 3.3E6 | 1.6E7 | nd | 5.7E6 | 1.2E7 | 3.1E7 |
| **PUA-Cal-HAD** | 1.9E7 | 8.3E6 | 5.6E6 | 5.4E6 | 1.8E6 | 6.7E6 | nd | 2.1E6 | 1.8E7 | 3.1E7 |
| | 1.5E7 | 1.3E7 | 4.8E6 | 6.7E6 | 8.7E5 | 8.2E6 | nd | 5.1E6 | 5.6E7 | 5.1E7 |
| | 1.7E7 | 1.3E7 | 4.4E6 | 5.6E6 | 1.2E6 | 8.2E6 | nd | 5.1E6 | 2.6E7 | 5.3E7 |
| **YFP T2 MOI 5** | 2.3E7 | 2.7E7 | 6.1E6 | 6.6E6 | 1.7E6 | 5.3E6 | nd | 1.2E7 | 2.9E7 | 2.8E7 |
| | 2.0E7 | 2.6E7 | 6.7E6 | 3.4E6 | 1.5E6 | 3.7E6 | nd | 8.1E6 | 2.4E7 | 2.6E7 |
| | 3.2E7 | 3.4E7 | 5.4E7 | 2.6E6 | 1.5E6 | 4.3E6 | nd | 3.8E6 | 2.8E7 | 2.8E7 |
| **PUA-Cal-HAD T2 MOI 5** | 2.9E6 | 3.2E6 | 5.7E5 | 1.2E6 | 1.7E6 | 4.8E6 | nd | 5.7E5 | 6.6E6 | 1.4E7 |
| | 5.8E6 | 5.0E6 | 7.2E5 | 2.1E6 | 2.7E5 | 7.7E6 | nd | 7.6E5 | 4.6E7 | 4.3E7 |
| | 8.4E6 | 6.4E5 | 1.3E6 | 2.3E6 | 4.1E5 | 5.0E6 | nd | 1.4E6 | 1.2E7 | 4.5E7 |

**Table S3** T2 escape mutants of PUA-Cal-HAD

| Phage | Gene | Locus_tag | Location | Mutation | Effect |
|---|---|---|---|---|---|
| T2m1 | *denA* | KMC23_gp183 | 133567 | C>CGG | Frameshift at R67 |
| | baseplate hub subunit and tail length | KMC23_gp215 | 114043 | C>T | P173L |
| T2m2 | *denA* | KMC23_gp183 | 133503 | G>GCTTC | Frameshift at S88 |
| | baseplate hub subunit and tail length | KMC23_gp215 | 114043 | C>T | P173L |
| | head-tail adaptor Ad2 | KMC23_gp250 | 90235 | G>A | G205R |

**Table S4** List of strains and phages used in this study, Related to STAR Methods

| Name | Reference | Sequence |
|---|---|---|
| **Strain** | | |
| *E. coli* DH10B | New England Biolabs (Cat # C3019) | NZ_CP110018.1 |
| *E. coli* BL21-AI | Thermo Fisher (Cat # C607003) | NZ_CP047231.1 |
| *E. coli* BL21 (DE3) | New England Biolabs (Cat # C2527H) | CP001509.3 |
| *E. coli* ECOR28 | Fagenbank | DABREU010000000 |
| *E. coli* BW25113 | Fagenbank | CP009273 |
| *E. coli* BW25113 Δdcm | KEIO collection | N/A |
| *E. coli* BW25113 Δdam | KEIO collection | N/A |
| **Phage** | | |
| *Escherichia* phage λ-vir | Fagenbank | NC_001416 [a] |
| *Escherichia* phage T1 | Fagenbank | NC_005833.1 |
| *Escherichia* phage T2 | DSMZ | NC_054931.1 |
| *Escherichia* phage T3 | Fagenbank | NC_003298.1 |
| *Escherichia* phage T4 | Fagenbank | NC_000866.4 |
| *Escherichia* phage T5 | Fagenbank | AY543070.1 |
| *Escherichia* phage T6 | Fagenbank | MH550421.1 |
| *Escherichia* phage T7 | Fagenbank | NC_001604.1 |
| Bas3 | Alexander Harms Basel Collection [1] | MZ501087 |
| Bas5 | Alexander Harms Basel Collection | MZ501101 |
| Bas10 | Alexander Harms Basel Collection | MZ501077 |
| Bas12 | Alexander Harms Basel Collection | MZ501053 |
| Bas13 | Alexander Harms Basel Collection | MZ501092 |
| Bas14 | Alexander Harms Basel Collection | MZ501107 |
| Bas16 | Alexander Harms Basel Collection | MZ501070 |
| Bas22 | Alexander Harms Basel Collection | MZ501091 |
| Bas24 | Alexander Harms Basel Collection | MZ501104 |
| Bas43 | Alexander Harms Basel Collection | MZ501106 |
| Bas47 | Alexander Harms Basel Collection | MZ501047 |
| Bas64 | Alexander Harms Basel Collection | MZ501081 |
| Bas66 | Alexander Harms Basel Collection | N/A |
| Bas67 | Alexander Harms Basel Collection | MZ501064 |

[a] Mutation in *cI* [2]

**Table S5** List of plasmids used in this study, Related to STAR Methods

| Plasmid | Description | Promoter | Resistance | Cloning method |
|---|---|---|---|---|
| pUOS031 | pACYC-YFP | pBAD | Cm | Gibson assembly |
| pUOS247 | pACYC-PUA-Cal | native | Cm | Gibson assembly |
| pUOS248 | pACYC-PUA-Cal inducible | pBAD | Cm | Gibson assembly |
| pUOS249 | pACYC-PUA-Cal-HAD | native | Cm | Gibson assembly |
| pUOS250 | pACYC-PUA-Cal-HAD inducible | pBAD | Cm | Gibson assembly |
| pUOS252 | pACYC-PUA-Cal-HAD PUA-Cal R10A | native | Cm | Around-the-horn on pUOS249 |
| pUOS254 | pACYC-PUA-Cal-HAD PUA-Cal W35A | native | Cm | Gibson assembly |
| pUOS256 | pACYC-PUA-Cal-HAD PUA-Cal W37A/W38A | native | Cm | Gibson assembly |
| pUOS257 | pACYC-PUA-Cal-HAD PUA-Cal K40A | native | Cm | Gibson assembly |
| pUOS259 | pACYC-PUA-Cal-HAD PUA-Cal R99A | native | Cm | Around-the-horn on pUOS249 |
| pUOS260 | pACYC-PUA-Cal-HAD PUA-Cal Y100A/Y101A | native | Cm | Around-the-horn on pUOS249 |
| pUOS264 | pACYC-PUA-Cal-HAD PUA-Cal D207A/H209A | native | Cm | Around-the-horn on pUOS249 |
| pUOS265 | pACYC-PUA-Cal-HAD PUA-Cal K246A | native | Cm | Around-the-horn on pUOS249 |
| pUOS268 | pACYC-PUA-Cal-HAD PUA-Cal H293A/D294A | native | Cm | Around-the-horn on pUOS249 |
| pUOS269 | pACYC-PUA-Cal-HAD PUA-Cal N292A | native | Cm | Around-the-horn on pUOS249 |
| pUOS270 | pACYC-PUA-Cal-HAD PUA-Cal N367A | native | Cm | Around-the-horn on pUOS249 |
| pUOS271 | pACYC-PUA-Cal-HAD PUA-Cal H462A/H464A | native | Cm | Around-the-horn on pUOS249 |
| pUOS272 | pACYC-PUA-Cal-HAD HAD D7A/D9A | native | Cm | Around-the-horn on pUOS249 |
| pUOS273 | pACYC-PUA-Cal-HAD HAD W15A | native | Cm | Around-the-horn on pUOS249 |
| pUOS274 | pACYC-PUA-Cal-HAD HAD E56A | native | Cm | Around-the-horn on pUOS249 |
| pUOS275 | pACYC-PUA-Cal-HAD HAD R114A/D276A | native | Cm | Around-the-horn on pUOS249 |
| pUOS276 | pACYC-PUA-Cal-HAD HAD T122A | native | Cm | Around-the-horn on pUOS249 |
| pUOS278 | pACYC-PUA-Cal-HAD HAD T147A | native | Cm | Around-the-horn on pUOS249 |
| pUOS279 | pACYC-PUA-Cal-HAD HAD K180A | native | Cm | Around-the-horn on pUOS249 |
| pUOS280 | pACYC-PUA-Cal-HAD HAD G204A/D205A/D210A | native | Cm | Around-the-horn on pUOS249 |
| pUOS281 | pCOLA-Gam | pBAD | Kan | Gibson assembly |
| pUOS282 | pCOLA-Ocr | pBAD | Kan | Gibson assembly |
| pUOS284 | pCOLA-P56 | pBAD | Kan | Gibson assembly |
| pUOS285 | pCOLA-AcrIF18 | pBAD | Kan | Gibson assembly |
| pUOS286 | pCOLA-UGI | pBAD | Kan | Gibson assembly |
| pUOS287 | pCOLA-Gp44 | pBAD | Kan | Gibson assembly |
| pUOS288 | pCOLA-AcrIF2 | pBAD | Kan | Gibson assembly |
| pUOS290 | pCOLA-ArdA | pBAD | Kan | Gibson assembly |
| pUOS292 | pCOLA-T4-Arn | pBAD | Kan | Gibson assembly |
| pUOS293 | pCOLA-T2-Arn | pBAD | Kan | Gibson assembly |
| pUOS294 | pCOLA-T4-Arn-N70S | pBAD | Kan | Gibson assembly |
| pUOS295 | pCOLA-Gam anderson 1 | J23100 | Kan | Gibson assembly |

| | | | | |
|---|---|---|---|---|
| pUOS297 | pCOLA-Gam anderson 0.1 | J23114 | Kan | Gibson assembly |
| pET01 | pET21a-PUA-His6 | T7 | Amp | Gibson assembly |
| pYB01 | pYB100-HAD-His6 | T5 | Kan | Gibson assembly |
| pYB02 | pYB100-HAD-Flag | T5 | Kan | Gibson assembly |
| pYB03 | pYB100-T2-Arn-Flag | T5 | Kan | Gibson assembly |
| pYBO4 | pYB100-T4-Arn-Flag | T5 | Kan | Gibson assembly |

### References


1. Maffei, E., Shaidullina, A., Burkolter, M., Heyer, Y., Estermann, F., Druelle, V., Sauer, P., Willi, L., Michaelis, S., Hilbi, H., et al. (2021). Systematic exploration of Escherichia coli phage–host interactions with the BASEL phage collection. PLOS Biol *19*, e3001424. 10.1371/journal.pbio.3001424.
2. Meyer, J.R., Dobias, D.T., Weitz, J.S., Barrick, J.E., Quick, R.T., and Lenski, R.E. (2012). Repeatability and Contingency in the Evolution of a Key Innovation in Phage Lambda. Science *335*, 428–432. 10.1126/science.1214449.
